# Age, Sex, and Genetics Influence the Abundance of Infiltrating Immune Cells in Human Tissues

**DOI:** 10.1101/614305

**Authors:** Andrew R. Marderstein, Manik Uppal, Akanksha Verma, Bhavneet Bhinder, Jason Mezey, Andrew G. Clark, Olivier Elemento

## Abstract

Despite infiltrating immune cells playing an essential role in human disease and the patient response to treatment, the central mechanisms influencing variability in infiltration patterns are unclear. Using bulk RNA-seq data from 53 GTEx tissues, we applied cell-type deconvolution algorithms to evaluate the immune landscape across the healthy human body. We first performed a differential expression analysis of inflamed versus non-inflamed samples to identify essential pathways and regulators of infiltration. Next, we found 21 of 73 infiltration-related phenotypes to be associated with either age or sex (*FDR* < 0.1). Through our genetic analysis, we discovered 13 infiltration-related phenotypes have genome-wide significant associations (iQTLs) (*P* < 5.0 × 10^−8^), with a significant enrichment of tissue-specific expression quantitative trait loci in suggested iQTLs (*P* < 10^−5^). We highlight an association between neutrophil content in lung tissue and a variant near the *CUX1* transcription factor gene (*P* = 9.7 × 10^−11^), which has been previously linked to neutrophil infiltration, inflammatory mechanisms, and the regulation of several immune response genes. Together, our results identify key factors influencing inter-individual variability of specific tissue infiltration patterns, which could provide insights on therapeutic targets for shifting infiltration profiles to a more favorable one.

## Introduction

Human immune systems vary dramatically across individuals, yet the environmental and genetic determinants of this variability remain poorly characterized. Many studies have identified genetic components and environmental stimuli that alter immune cell composition in peripheral blood^1–6^. However, normal tissues and organs also consist of diverse cell types, including infiltrating immune cells. The variability in infiltration across individuals and between tissues has not been documented and the mechanisms enabling such variation in baseline infiltration have not been elucidated.

Identifying the genetic influences on specific patterns of infiltrating immune cells is crucial to understanding disease biology. Beyond further explaining heritable manifestations of infectious diseases and autoimmunity^1–9^, such efforts can further uncover the drivers of characteristic immune cell signatures in the tumor microenvironment that are prognostic for cancer progression and predictive of treatment response^10^. For example, response is improved in patients with T cell-inflamed tumors compared to T cell-depleted tumors among patients receiving immune checkpoint inhibitors targeting PD1 and CTLA4^10,11^ and among ovarian cancer patients receiving chemotherapy^12^. However, a complete mechanistic description underlying immune-rich and immune-poor tumor phenotypes remains elusive.

Recent advances in computational methods have allowed reliable inference of the heterogeneous cell types from gene expression data of a single population-level (bulk) tissue sample^13–15^. At the same time, large-scale sequencing efforts such as the GTEx project^16^ have enabled a detailed exploration of the links between genomic and transcriptomic variations across different tissues. Together, these cell-type estimation methods can be utilized in synergy with massive bulk sequenced data sets to infer cellular heterogeneity and achieve statistically well-powered associations that intrinsically drive the heterogeneity^17–19^.

In the present study, we aimed to evaluate the inherent immune infiltration landscape across healthy tissues in the human body and to determine the intrinsic factors contributing to the infiltration variability. Using bulk RNA-seq data from 53 distinct GTEx tissue types, we applied cell-type deconvolution algorithms to infer immune content and developed an analysis framework to leverage information across methods for association testing. We identified transcriptomic differences associated with extreme infiltration patterns by performing a differential expression analysis of the immune-rich and immune-depleted samples^20^. We discovered associations between donor characteristics such as age, sex, and germline genetic variants with infiltration variability. We find that these genetic determinants are enriched for an overlap with tissue-specific expression quantitative trait loci (eQTLs). Additionally, such genetic variants can serve as leading candidates for understanding the basic biology behind infiltrating immune cell patterns.

## Results

### Robust estimation of immune cell types in bulk RNA-seq profiles

To describe immune content from bulk RNA-seq samples, we used two central algorithms: xCell^13^ and CIBERSORT^14^. xCell relies on a modification of single sample gene-set enrichment analysis to estimate cell type scores, while CIBERSORT employs a linear support vector regression model. The default reference signatures allow deconvolution of 64 immune and stroma cell types for xCell and 22 immune cell types for CIBERSORT. CIBERSORT also calculates a scaling factor that measures the degree of infiltration. We refer to the relative proportions from CIBERSORT as “CIBERSORT-Relative” and the product of the relative proportions with the scaling factor as “CIBERSORT-Absolute”. We estimate three scores for each cell type to describe the immune content from the gene expression data for each tissue in each individual: xCell, CIBERSORT-Relative, and CIBERSORT-Absolute scores.

We first hypothesized that the relative and absolute scores from CIBERSORT encapsulated different aspects of the single-cell deconvolution. While “CIBERSORT-Absolute” simultaneously quantifies a degree of immune infiltration, “CIBERSORT-Relative” is focused on capturing compositional changes in the immune content (Supplementary Note). We simulated synthetic mixes composed of bulk tissue “spiked” in silico with CD4+ T cells and CD8+ T cells (see **Methods**). We correlated the known amount of CD4+ and CD8+ T cell infiltration in these mixtures with estimated deconvolution scores under a “tissue” scenario and an “immune cell” scenario. In the “immune cell” scenario, we let the true infiltration be the proportion of each cell type to the total immune content. In the “tissue” scenario, the true infiltration amount is the proportion of each cell type to the entire sample. As expected, we found that CIBERSORT-Relative to more accurately estimate the infiltration amounts of the in silico mixtures than CIBERSORT-Absolute in the “immune cell” scenario, while the reverse was true in the “tissue” scenario (Supplementary Table 1). However, in both scenarios, the CIBERSORT method resulted in strong correlations between deconvolution scores and the true amount of infiltration (*r* = 0.64-0.89).

We also compared the CIBERSORT performance to xCell. Since xCell is an enrichment-based algorithm, not a deconvolution algorithm, it is not recommended for comparing scores between cell types. As a result, xCell scores correlated well in the “tissue” scenario (*r* = 0.90 with CD4+ T cells, *r* = 0.96 with CD8+ T cells) but worse in the “immune cell” scenario (*r* = 0.65 with CD4+ T cells, *r* = 0.45 with CD8+ T cells), even after normalization (Supplementary Table 2). We also found that xCell had imperfect correlation with CIBERSORT-Absolute scores (CD8: *r* = 0.58, CD4: *r* = 0.86). Therefore, these results indicate that each method provides interesting information to be exploited in downstream analysis.

### Evaluating infiltration across human tissues by using deconvolution

Our first objective was to deconvolute cellular heterogeneity from bulk RNA-seq GTEx samples and examine the immune infiltration profile. The v7 GTEx release consists of 11,688 samples, spanning 53 different tissues/sample types from 714 donors^16^. We performed comprehensive deconvolution of these samples using our three methods (xCell, CIBERSORT-Relative, and CIBERSORT-Absolute) and focused most of our analyses on CD4+ T cells, CD8+ T cells, macrophages, and neutrophils (Figure 1a; Supplementary Note).

To obtain a global overview of heterogeneity in immune composition, we performed hierarchical clustering of the tissues based on the medians of each immune cell type (see **Methods**) (Figure 1b). Across the 3 methods, many of the nearest-neighbor pairings were consistent and recapitulated relationships between tissues that share high degrees of histologic similarity and immune infiltration (Supplementary Note; Supplementary Figures 4-5). In our analysis of macrophage content across tissues, we found the highest scores (CIBERSORT-Absolute) to be in lung, spleen, and adipose tissue (Figure 1c). High levels of macrophages, especially uncommitted M0 and anti-inflammatory M2 macrophages, were found in lung tissues (Figure 1b). We discuss other interesting observations comparing infiltration patterns between tissues in the Supplementary Note (Supplementary Figures 6-9).

Importantly, we found large variability between different individuals in a single tissue type. Many tissues featured a majority of samples with trace immune enrichment, but these tissues also contain several samples with significantly higher estimated immune content (Supplementary Figures 6-9). Interestingly, t-SNE visualizations of estimated immune content within a tissue type did not reveal distinct clusters of samples (Supplementary Figure 10). Therefore, it appears that healthy individuals have highly variable infiltration patterns, suggesting that the differences could be driven by a range of genetic and non-genetic factors (such as age, sex, or environmental exposures). We aimed to identify transcriptomic associations with infiltration through a differential expression analysis of grouped immune-enriched (inflamed) vs immune-depleted (non-inflamed) tissue samples, and we aimed to identify genetic effects through a genome-wide association study (GWAS) analysis of immune content. Through a carefully designed filtering procedure, we focus our analysis on a limited set of 73 infiltration phenotypes that represent immune cell type scores in a tissue (tissue-by-cell type pairs) (see **Methods**).

Lastly, we were interested in whether infiltration signatures explain substantial variance of gene expression calculated in bulk assays. We performed a principal component analysis of the processed gene expression matrix within each tissue, before assessing the pairwise relationship between the first four principal components and the 73 infiltration phenotypes using CIBERSORT-Absolute scores. Even after a Benjamini-Hochberg false discovery rate (FDR) correction^21^, we found that all but one infiltration phenotype was significantly correlated with at least one principal component (FDR < 0.1), indicating that bulk gene expression measurements are significantly confounded by cellular heterogeneity (Supplementary Table 9).

**Figure 1:**
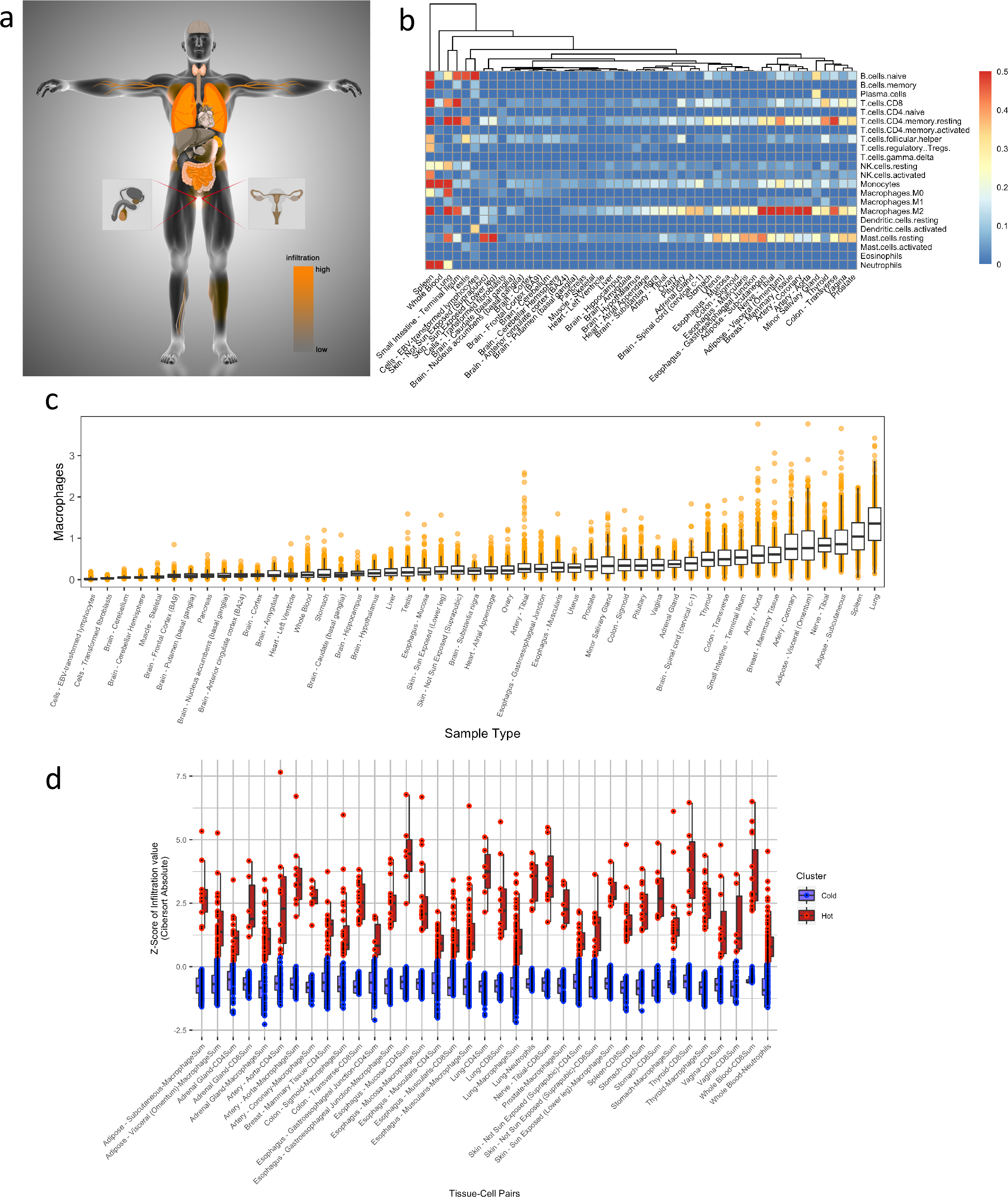
Immune content across the human body. (a) Human body overlaid with GTEx tissues colored by the degree of immune infiltration, as estimated by the scaling factor from CIBERSORT-Absolute. (b) Hierarchical clustering of GTEx tissues according to immune content (estimated by CIBERSORT-Absolute). Heatmap displays cell type median scores, with upper bound set to 0.5. (c) Macrophage content across tissues. Scores estimated by CIBERSORT-Absolute and sorted. (d) Differential immune content between “hot” and “cold” clusters of the 43 infiltration phenotypes for which DEGs could be identified.

### Identification and characterization of extreme infiltrating immune cell patterns

We then searched for genes preferentially expressed in immune cell type rich (hot) versus immune cell type depleted (cold) cases. For each tissue type, we used consensus k-means clustering to identify hot and cold cases for each cell type (Supplementary Table 3; Figure 1d). We then performed a differential gene expression analyses between corresponding hot and cold cases for each cell type. These identified differentially expressed genes (DEGs) may partially explain tissue-specific heterogeneity in baseline immune infiltration.

Overall, we identified DEGs for 43 of the infiltration phenotypes which passed our statistical thresholds (log FC >= 2.0, FDR < 0.01; see **Methods**). Across the 10 CD8+ T cell, 9 CD4+ T cell, 21 macrophage, and 3 neutrophil phenotypes tested, we expected and found that the most common DEGs consisted of well-known markers of the corresponding immune cell types (Supplementary Table 5). For example, the most consistent DEGs across macrophage-hot clusters are macrophage markers utilized by the xCell and CIBERSORT algorithms for estimating macrophage content: *C1QB* (18/21 tissues), *VSIG4* (17/21), *MARCO* (17/21), and *CD163* (16/21). Interestingly, the most common DEGs not present in the deconvolution reference gene sets includes *C1QC* (16/21) and *FCGR3A* (16/21), which correspond to complement component and immunoglobulin Fc receptor, and have well-characterized roles in opsonization ^22,23^. We then used the DEGs and Ingenuity Pathway Analysis (IPA) to identify dysregulated pathways, discover key upstream regulators, and find central disease and function ontologies (Supplementary Table 6, 7, 8). In our macrophage phenotypes, the most commonly dysregulated pathways were *TREM1* signaling, which is an amplifier of macrophages inflammation^24^, and antigen-presenting cell maturation (Supplementary Table 6). The most frequent upstream regulator predicted to be activated by IPA in the macrophage-hot clusters was *TGM2*, while *TFRC* (transferrin receptor) was the most commonly inhibited as predicted by IPA (Supplementary Table 7). The latter finding may be linked to the role of macrophages in sequestering iron during inflammatory states^25^. Disease and function ontologies indicated that the most commonly activated pathways in macrophage-hot samples were associated with leukocyte and lymphocyte migration (Supplementary Table 8). We describe the results across the T cell and neutrophil phenotypes in Supplementary Note).

Finally, we used our immune-hot clusters (eg. macrophage-hot) to examine whether individuals with inflammation in one tissue type may also exhibit similar inflammation in their other tissue types. Here, we report inconsistent inflammation patterns across distinct tissue types within the same individual (mode = 1 tissue per individual for each cell type) (Supplementary Figure 11). Therefore, we reflected that infiltration patterns are likely tissue-specific, rather than widespread.

### Association of age and sex with immune infiltration

We next aimed to examine whether there are any associations of age or sex with immune infiltration. We adopted a multiple regression approach to measure age and sex effects across all infiltration phenotypes. This was repeated for each of the deconvolution procedures, and we merged p-values across all three by using Empirical Brown’s method^26^ (see **Methods**).

We observed that 21 of 73 infiltration phenotypes were significantly associated with either age or sex (Table 1). While similar numbers of sex and age associations were identified (phenotypes with *FDR* < 0.1: 12 for age, 13 for sex), we found the most significant associations were increased T cell content in female breast tissue compared to male (Sex-CD8+ T cell association: *P* = 5.4 × 10^−34^ (Figure 2a); sex-CD4+ T cell associations: *P* = 3.2 × 10^−9^). In female samples, we observed significant heterogeneity, with several samples having no CD8+ T cell content detected and others having high predicted CD8+ T cell content (Figure 2a). The distinct contrast between female and male breast tissue could drive these immune differences, with male tissue predominantly lacking the lobular elements^27^ and temporal changes of females that are associated with T cells^28^ (Supplementary Note). However, the key drivers of this variability pattern remain unclear.

The most significant association with age is CD4+ T cells in tibial artery tissue (*P* = 1.3 × 10^−9^) (Figure 2b). We noted that 4/5 tested infiltration phenotypes from tibia area (exception: macrophage content in tibial nerve samples) had increased immune cell scores with age. In comparison, only 1/6 phenotypes in artery tissue from other body areas showed significant infiltration patterns with age (CD4+ T cells content in arterial aorta samples). Again, the reasons for these localized associations are unclear.

**Table 1:** Significant associations with (a) age and (b) sex. Significance indicates that FDR-adjusted p-values are below 0.1. Raw and adjusted p-values are shown. +/− indicates an increase/decrease in older age samples, or an increase/decrease in females compared to males samples.

| Tissue | Cell Type | Effect Direction | P-value | Adj P-value |
| --- | --- | --- | --- | --- |
| Artery - Tibial | CD4+ T cells | + | 1.3e-09 | 9.7e-08 |
| Artery - Tibial | Macrophages | + | 3.9e-05 | 7.2e-04 |
| Adipose - Visceral (Omentum) | CD8+ T cells | + | 1.7e-03 | 1.8e-02 |
| Breast - Mammary Tissue | Macrophages | + | 5.2e-04 | 7.5e-03 |
| Thyroid | CD8+ T cells | + | 6.1e-04 | 7.5e-03 |
| Lung | CD4+ T cells | + | 4.1e-03 | 3.2e-02 |
| Esophagus - Muscularis | CD4+ T cells | - | 3.5e-03 | 3.2e-02 |
| Small Intestine - Terminal Ileum | CD8+ T cells | - | 4.3e-03 | 3.2e-02 |
| Nerve - Tibial | CD8+ T cells | + | 3.0e-06 | 1.1e-04 |
| Nerve - Tibial | CD4+ T cells | + | 2.3e-05 | 5.7e-04 |
| Whole Blood | Neutrophils | - | 7.0e-03 | 4.7e-02 |
| Artery - Aorta | CD4+ T cells | + | 8.9e-03 | 5.5e-02 |

+/- indicates increase/decrease with older age b
| Tissue | Cell Type | Effect Direction | P-value | Adj P-value |
| --- | --- | --- | --- | --- |
| Adipose - Subcutaneous | Macrophages | + | 8.5e-06 | 9.2e-05 |
| Breast - Mammary Tissue | CD8+ T cells | + | 5.4e-34 | 3.5e-32 |
| Breast - Mammary Tissue | CD4+ T cells | + | 3.2e-09 | 1.0e-07 |
| Breast - Mammary Tissue | Macrophages | - | 1.4e-06 | 2.1e-05 |
| Thyroid | CD8+ T cells | + | 1.6e-06 | 2.1e-05 |
| Thyroid | CD4+ T cells | + | 1.1e-06 | 2.1e-05 |
| Thyroid | Macrophages | + | 4.7e-03 | 3.1e-02 |
| Lung | CD4+ T cells | + | 1.4e-02 | 7.1e-02 |
| Esophagus - Muscularis | CD4+ T cells | + | 1.5e-03 | 1.2e-02 |
| Esophagus - Muscularis | Macrophages | + | 5.4e-03 | 3.2e-02 |
| Whole Blood | Neutrophils | + | 3.7e-03 | 2.7e-02 |
| Artery - Aorta | CD8+ T cells | - | 7.0e-03 | 3.8e-02 |
| Pituitary | CD8+ T cells | - | 1.2e-03 | 1.1e-02 |

+/- indicates increase/decrease in females

### Association of genetic variants with infiltrating immune cells

We next searched for particular inherited genetic variants that could influence the variability of infiltration patterns. We refer to germline single nucleotide polymorphisms (SNPs) associated with the infiltration of a cell type in a tissue as infiltration quantitative trait loci (iQTLs). Using the Empirical Brown’s testing framework across 73 infiltration phenotypes, we discovered 13 infiltration phenotypes with at least one genome-wide significant iQTL (*P* < 5.0 × 10^−8^) and 2 phenotypes with at least one study-wide significant iQTL (*P* < 6.8 × 10^−10^) (Table 2) (see **Methods**).

The most significant iQTL we identified was an association between rs77155650 and neutrophil content in lung samples (*P* = 9.7 × 10^−11^) (Figure 2c-d) (Supplementary Note). The variant lies 76 kb from the transcription factor *CUX1*, in a potentially active regulatory region of the non-coding genome that overlaps enhancer histone marks across most cell types and overlaps promotor histone marks and DNAse sites in some cell types^29,30^. The DNA binding activity of the *CUX1* protein product has been previously linked with neutrophil infiltration^31,32^ and the regulation of multiple immune response genes^33^ (*F2RL1*^34,35^, *IL1A*, *MMP10*^36–38^, and *COX2*^34,39^) (Supplementary Note). In the GTEx lung samples, we found that *CUX1* expression in the lung samples correlated significantly with neutrophil infiltration (*P* = 4.5 × 10^−4^) (Figure 2e). Potentially, the rs77155650 polymorphism could interact with CUX-1 DNA binding and the regulation of immune response genes in inflammation processes.

The second study-wide significant iQTL we discovered was an association between rs116827016 and macrophage infiltration in tibial artery tissue (*P* = 3.86 × 10^−10^) (Table 2; Figure 2f-g) (Supplementary Note). This SNP has not been identified as an eQTL in any tissue within the GTEx consortium analyses^16^, but has been commonly associated as an eQTL for *KCTD10* expression in whole blood (eQTLGen meta-analysis p-value = 5.7 × 10^−63^)^40^. We discovered a significant correlation between the *KCTD10* expression in the GTEx tibial artery samples with the macrophage phenotype (*P* = 3.2 × 10^−16^) (Figure 2h). We analyzed *KCTD10* expression across tissues and found that expression is highest in tibial artery samples (Supplementary Figure 13), and the expression-infiltration association is driven by increased macrophages in low *KCTD10*-expressed samples (Figure 2h). Functional studies of *KCTD10* and its paralog *TNFAIP1* have linked both to inflammation-associated angiogenesis^41,42^, so its possible that the rs116827016 haplotype alters immune response within the vascular system through changes in *KCTD10* expression (Supplementary Note).

Variants already associated with gene expression allow inference into functional roles. Thus, we were next interested in identifying whether there were expression quantitative trait loci (eQTLs) from the GTEx consortium analysis^16^ that were also iQTLs (ieQTLs). We found that 2 infiltration phenotypes had ieQTLs surpassing genome-wide significance (*P* < 5.0 × 10^−8^) (Table 2).

Our third most significant iQTL association (and most significant ieQTL association), the rs11883564 locus with CD4+ T cells content in sun exposed skin tissues (*P* = 4.2 × 10^−9^), overlapped with a significant association for *STAM2* gene expression^16^ (Table 2; Figure 2i-j). *STAM2* is essential for T-cell development^43^ and the rs11883564 haplotype has been associated with alopecia areata response to chemotherapy in breast cancer patients^44^. Alopecia areata is an autoimmune disease characterized by hair loss from infiltrating T cells^45^. We also discovered significant correlation between *STAM2* gene expression and the CD4+ T cell signature in skin tissue (p = 7.2 × 10^−9^; Figure 2k). Genetic network analysis of *STAM2* revealed an integral role in immune regulation pathways (Supplementary Figure 14, Supplementary Note, see **Methods**).

**Figure 2:**
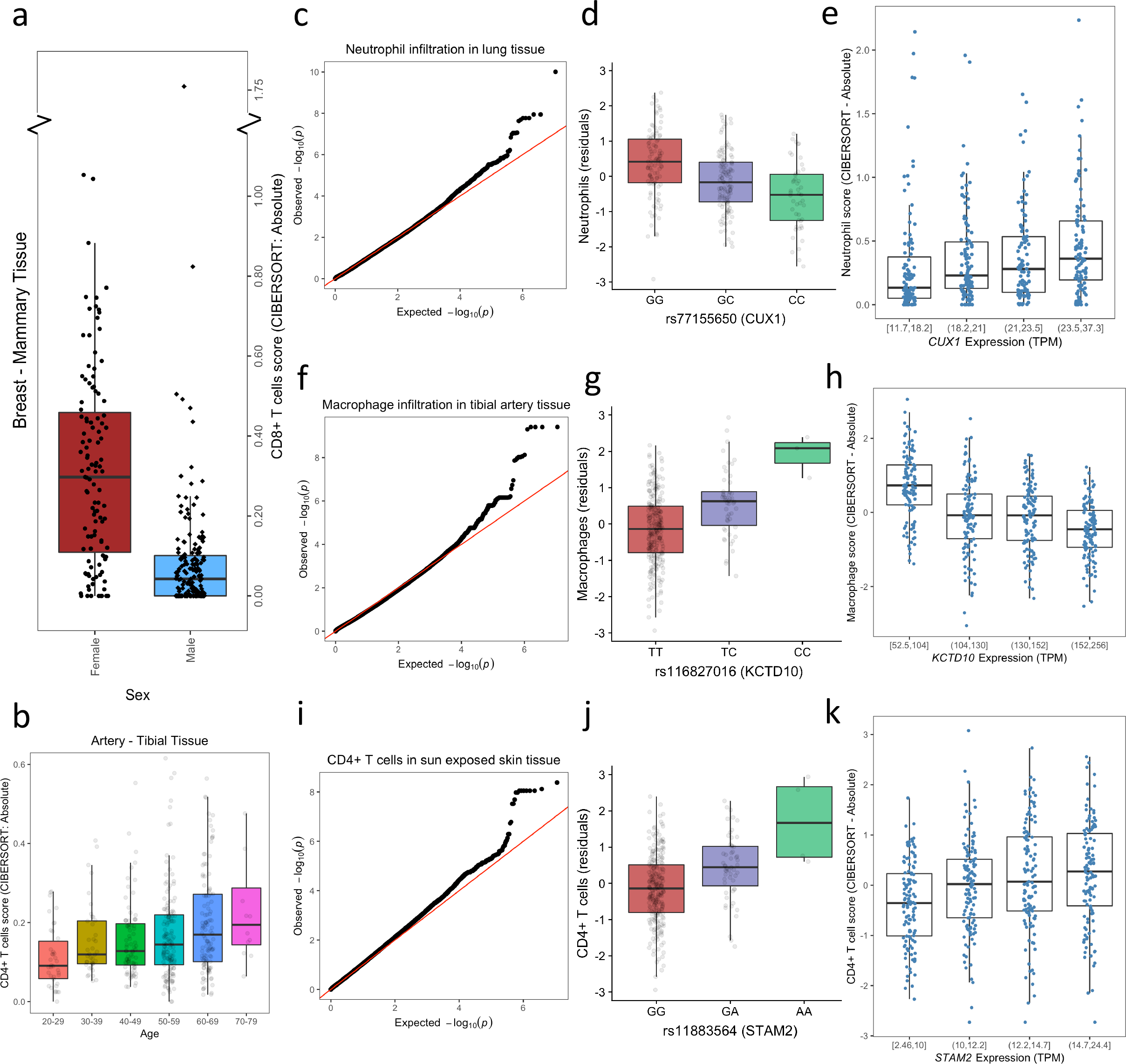
Significant associations with infiltrating immune cell patterns. (a) shows sex association with CD8+ T cell content in breast tissue samples. (b) shows age association with CD4+ T cell content in tibial artery samples. (c-e) relates to the genetic analysis of neutrophil content in lung tissue samples, (f-h) is macrophage content in tibial artery tissue, and (i-k) concerns the CD4+ T cell content in sun exposed skin tissue. The leftmost of these plots are genome-wide QQ-plots of Empirical Brown’s p-values, the middle plots reflect genotype-phenotype association plots (using CIBERSORT – Absolute residuals), and the rightmost plots are eGene expression-phenotype association plots of the significant iQTL (using estimated scores from CIBERSORT – Absolute), split into quartiles of gene expresson.

### Downstream analysis of genetic results

Next, we systematically tested for an enrichment of tissue-specific expression QTLs in the most significant SNPs from our genetic analysis. We first relaxed our iQTL threshold to the GWAS catalog cut-off by querying all genetic associations with p < 10^−5^ to represent our “top hits” and developed two complementary approaches for testing over- or underrepresentation of tissue-specific eQTLs (see **Methods**). Both approaches converged in showing significant enrichment of tissue-specific eQTLs in the top hits for many phenotypes (Figure 3; Supplementary Figure 15) (Supplementary Note). We note that directionality is unclear when SNPs are both iQTLs and eQTLs. These SNPs could independently alter immune content and expression levels, but it is also likely that differences in expression levels drive changes in infiltration patterns. Similarly, it is likely that cellular heterogeneity differences (from infiltration effects) underlie many eQTL associations.

To attempt to draw functional conclusions from our genetic results, we used the gene expression associations from our ieQTLs. We constructed a GeneMania network^46^ by forming a list of ieGenes (genes whose expression is associated with the variant, as determined from GTEx analysis^16^) from ieQTLs with our relaxed iQTL threshold p < 10^−5^ (GWAS catalog cut-off). We queried 85 genes, building a network of 115 total genes with 15 functional attributes and 30 additional genes (Supplementary Figure 16). We found that this network was enriched for pyrimidine biosynthesis and DNA repair functions (Supplementary Table 12). These functions help maintain healthy DNA, which is crucial for normal function and cancer avoidance. We discovered that the most interconnected added genes were involved in pyrimidine biosynthesis, transcriptional mechanisms, epigenetic remodeling, and immune-related disorders (Supplementary Note). Lastly, GeneMania identified a 9.55% weighting enrichment between network nodes to the Gasdermin protein domain, as collected in InterPro^47^. Gasdermin is required for recognizing foreign material and pathogens to induce IL-1 for recruiting immune cells^48,49^. In summary, our GeneMania network highlights central mechanisms controlling the immune environment.

Lastly, we analyzed whether iQTLs commonly displayed pleiotropic effects. We found that almost all iQTLs were associated with only a single cell type in a single tissue type under a p < 10^−5^ threshold (Supplementary Table 13). We also found that 3/176 ieGenes from ieQTLs (using the relaxed p < 10^−5^ threshold) were phenotype-specific (Supplementary Table 14; Supplementary Figure 17).

**Table 2:** Genome-wide significant variant associations (*P* < 5 × 10^−8^) for the 73 tested infiltration phenotypes. For each phenotype, only the most significant SNP is listed. The association *q*-values and whether the significant SNP is a GTEx eQTL in that tissue is also listed. Chromosome and position are reference genome build 37.

| Tissue | Cell | RS ID | Chromosome | Position | P-value | q-value | eQTL? |
| --- | --- | --- | --- | --- | --- | --- | --- |
| Lung | Neutrophils | rs77155650 | 7 | 101382863 | 9.76E-11 | 0.0003 | FALSE |
| Artery - Tibial | Macrophages | rs116827016 | 12 | 109787397 | 3.86E-10 | 0.0003 | FALSE |
| Skin - Sun Exposed (Lower leg) | CD4+ T cells | rs11883564 | 2 | 153016576 | 4.15E-09 | 0.0040 | TRUE |
| Esophagus – Gastro. Junction | CD4+ T cells | rs9847949 | 3 | 24599271 | 6.54E-09 | 0.0157 | FALSE |
| Vagina | CD8+ T cells | rs67160095 | 5 | 16047889 | 9.21E-09 | 0.0226 | FALSE |
| Esophagus - Mucosa | Macrophages | rs12631070 | 3 | 1518915 | 1.48E-08 | 0.0280 | FALSE |
| Spleen | Neutrophils | rs10956711 | 8 | 134621178 | 2.07E-08 | 0.0395 | FALSE |
| Nerve - Tibial | CD4+ T cells | rs9422297 | 10 | 1257108276 | 2.08E-08 | 0.1046 | FALSE |
| Thyroid | Macrophages | rs7277675 | 21 | 42637222 | 2.51E-08 | 0.0494 | TRUE |
| Skin - Sun Exposed (Lower leg) | Macrophages | rs4118372 | 3 | 189287063 | 2.56E-08 | 0.0318 | FALSE |
| Artery - Aorta | CD8+ T cells | rs34573417 | 10 | 9576753 | 2.85E-08 | 0.0494 | FALSE |
| Colon - Transverse | Macrophages | N/A | 10 | 3629677 | 2.94E-08 | 0.0776 | FALSE |
| Prostate | CD8+ T cells | rs12603004 | 17 | 13712941 | 4.26E-08 | 0.0435 | FALSE |

**Figure 3:**
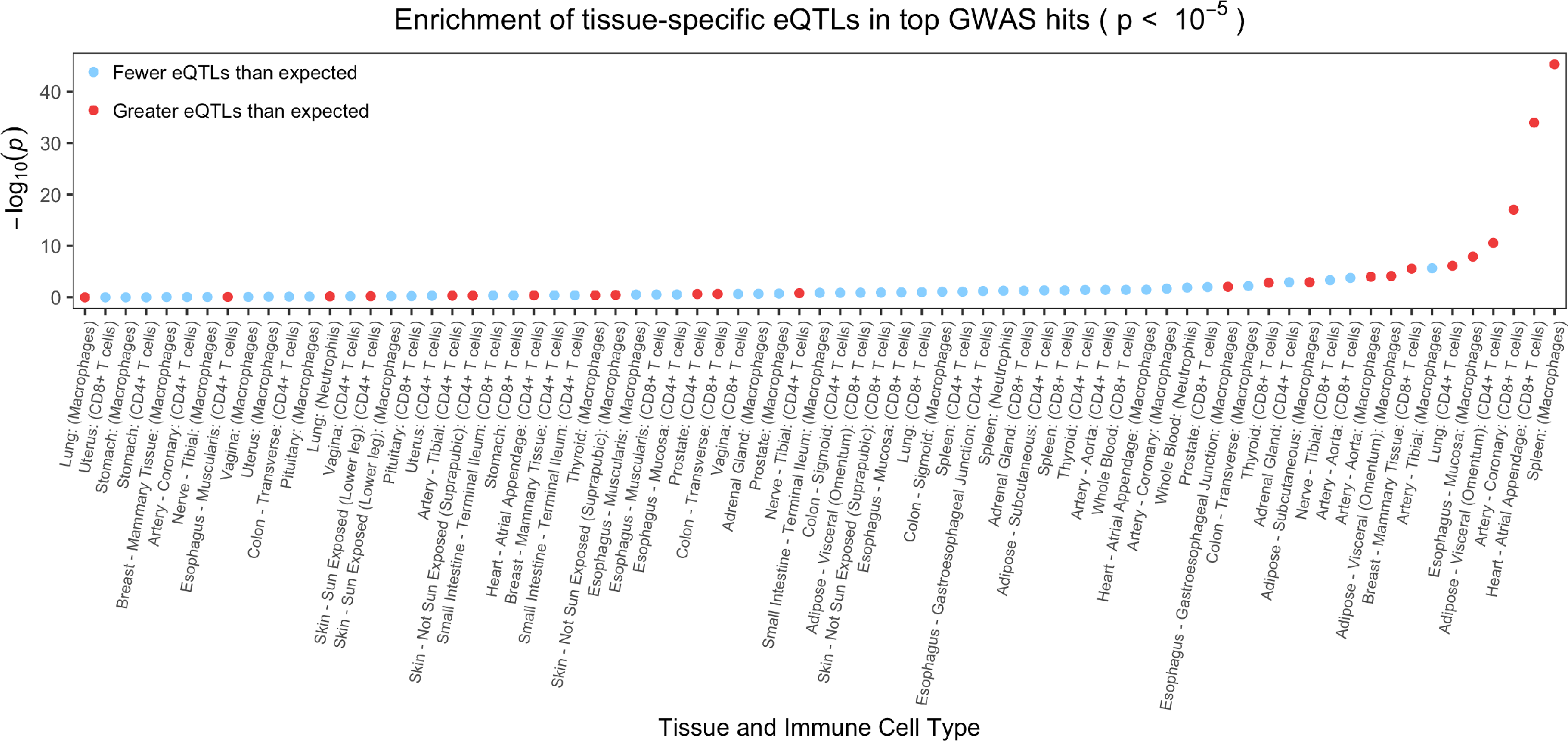
eQTL enrichment in iQTLs. Test 1 chi-square p-values across all infiltration phenotypes, colored by whether eQTLs are over-represented (red) or under-represented (blue) in the iQTLs.

## Discussion

The GTEx consortium project enabled an analysis of transcriptomic variation across diverse human tissues, and the discovery of association between that variation and genetic polymorphisms. With the development of computational algorithms that can deconvolute the cellular heterogeneity underlying bulk RNA-seq data, the GTEx data sets could be utilized to evaluate the baseline immune landscape across the human body. This quantification of immune cells from bulk RNA-seq should become standard in many bulk RNA-seq analyses. Additionally, we observed that population-level gene expression values are strongly affected by infiltrating immune cells through a principal component-based correlation analysis, implying that measures of infiltration should be considered in downstream analyses of bulk expression data.

We also developed frameworks to leverage information across multiple cell-type estimation methods and capture differences in deconvolution methods. An individual cell-type estimation method has imperfect correlation with the true scores and other computational methods, likely due to the selected markers and cell types in the reference set inducing certain biases. Thus, we increased our confidence in downstream analyses by incorporating results from multiple deconvolution algorithms, and demonstrated convergence on plausible, significant results while maintaining low false positive rates in large genomic analyses.

We also note that in the clinic, the relative ratio of CD4:CD8+ T cells is a blood test marker to monitor the health of the immune system^50^, which can be better captured by relative estimates of infiltration. Our demonstration of heterogeneous infiltration profiles across tissues suggests that deconvolution methods can potentially be used to derive an expanded set of biomarkers to assess immunologic health across a variety of organs.

Importantly, we demonstrated substantial variability in immune content across individuals within a number of different tissue types. Inspired by efforts to characterize the heterogeneity of tumor immune landscapes, we engaged in an endeavor to discretize clusters of individuals based on their tissue infiltration^20,51,52^. We were able to successfully show strong separability of immune content in hot clusters compared to cold ones across most phenotypes, and used these clusters to discover that the strongest DEGs revealed essential functions, pathways, and upstream regulators. Furthermore, we identified genetic loci associated with immune infiltration that could pose as future clinical markers to guide patient stratification. In cancer, the infiltration profile may be driven by not only new somatic mutations but also pre-existing germline variants. Since the immune signature in the tumor microenvironment is highly correlated with the response to treatments such as immunotherapy, germline variants could enhance predictive modeling of response and reveal novel therapeutic targets for shifting infiltration profiles to a more favorable one. Previous studies demonstrated that cancer cells maintain chromatin structure from the tissue-of-origin, so it is possible that germline iQTLs have conserved infiltration effects in the cancer cells^53^. If this were the case, then functional experiments could be a promising avenue for developing medicines to shift infiltration patterns. Overall, understanding a personalized baseline immune response from genetics would enhance the interpretation of immune presence in the tumor microenvironment.

An important area of future research is to test associations between somatic mutation burden and immune infiltration estimates in healthy tissues. Recent research performed using the GTEx database has shown that genetically distinct non-cancerous subclonal populations may arise in healthy tissues, akin to the phenomenon of clonal hematopoiesis of indeterminate potential in blood cells^54^. We saw a significant increase of T cell content in female breast and thyroid tissues compared to males. Immune response may correlate with somatic mutation detection, and it would be interesting to evaluate whether differential immune content between males and females is driven by sex-differential somatic clonal mutations which could eventually promote cancer. In parallel, the epidemiological differences between males and females in developing breast and thyroid cancer is striking: of breast cancer, there are 100 times more cases in females compared to males, and of thyroid cancer, there are nearly 3 times as many cases in females compared to males^55^. Further investigation could provide improved understanding of the increased female disease incidence.

Lastly, we note that superior computational algorithms for cell-type estimation are still needed, as well as larger and better annotated data sets. The algorithms we used are limited to inference of single-cell compositional information and do not infer a cell’s molecular signatures, where dysregulation may be even more informative^56^. While single-cell sequencing can provide a more intricate and accurate perspective of the infiltrating immune cells, single-cell studies have not been scaled large enough to understand the genetic basis of infiltration patterns. Even with bulk sequencing, the sample sizes examined in our study are limited and must be expanded. We limited our statistical genetic analysis to tissues having greater than 70 samples with matched genotype and phenotype information, and the maximum tissue type had 361 samples. At this sample size, we can only detect the largest of genetic effects. If infiltration is a widely polygenic trait, then increased sample size is necessary to dissect the genetic architecture of inter-individual differences in infiltration. Similarly, larger sample sizes will allow potential detection of tissue-specific immunomodulatory genes in our hot-cold analysis. Finally, it would enable improved assessment of infiltration pleiotropy. In our study, our identified genetic variants were rarely associated with multiple immune infiltration phenotypes. This implies that the genetics of infiltration differs depending on the tissue of interest and the expression patterns in that tissue, and that down-regulation of one tissue’s key functional genes within another tissue could create a completely separate genetic variation network that leads to infiltration. However, a larger dataset is necessary to ascertain how tissue-specific the genomics of infiltration patterns are.

## Methods

### GTEx data

Processed gene expression profiles from the GTEx v7 data release were downloaded from the GTEx data portal. Genotype data and raw fastq reads are from the GTEx v6 release, and were downloaded from dbGAP.

### Deconvolution of bulk RNA-seq profiles

To deconvolute bulk RNA-seq profiles into single-cell scores, we used CIBERSORT-Relative, CIBERSORT-Absolute, and xCell. CIBERSORT outputs were generated using 1000 permutations and quantile normalization disabled. Since xCell scores are generated by assessing relative variability between all input samples, the scores were generated separately for each tissue type. This allows better sensitivity to within-tissue cellular variability without substantial confounding from between-tissue variability, which was proven by our testing of xCell on synthetic mixes (Supplementary Table 2). Each cell type by tissue type combination is considered an infiltration phenotype.

### Simulating “immune-spiked” synthetic mixes

To generate “immune-spiked” synthetic mixes, we hand-selected one sigmoid colon GTEx sample (GTEX-XXEK-1826-SM-4BRVC) and one sun-exposed skin GTEx sample (GTEX-WFON-2126-SM-3LK7O) from the v6 release. Both these samples were identified by applying CIBERSORT-Absolute to all GTEx samples and identifying samples with the lowest detected presence of infiltrated immune cells (high CIBERSORT p-values, low cell scores). Using 5 different CD4+ T cell references and 5 different CD8+ T cell references (Supplementary Table 15 for SRA), we designed 90 synthetic mixes which contained 80-95% reads sampled from one of the GTEx samples and 5-20% of the reads sampled from the T cell samples. There were four different simulation types: (1) only CD4+ T cells infiltration as 5-20% of the sample, (2) only CD8+ T cells infiltration as 5-20% of the sample, (3) both CD4+ and CD8+ T cells infiltration in equal proportions as 5-20% of the sample, and (4) CD4+ and CD8+ T cells infiltration but in unequal proportions (2:3 and 1:4 ratios as 5-20% of the sample). Half the simulations were created using 1 CD4+/CD8+ reference and half with 5 CD4+/CD8+ references (cellular heterogeneity versus no heterogeneity). Both the colon and skin samples represented half the simulations, and the skin and colon samples were not part of any of the same synthetic mixes. The different immune samples used and their relative proportions across the 90 synthetic mixtures generated is outlined in Supplementary Table 15.

The bulk tissue and immune samples were aligned to the GRCh38 reference genome using *STAR*^57^ and sorted with *samtools*^58^. The number of reads in each sample were measured using *samtools idxstats*, then downsampled to the desired library size using *samtools view* with the -*s* flag and the specified percentage of total reads. Next, the resulting bam files containing the downsampled bulk and immune reads were merged using *bamtools merge* to create a single synthetic mixture bam file^59^.

### Generating TPM gene measurements from the synthetic mixes

RNAseq samples were quantified with the Gencode gene annotation reference (V22 release). Aligned reads were then quantified for gene expression in terms of TPM and FPKM using StringTie^60^.

### Empirical evaluation of CIBERSORT relative vs absolute outputs

To empirically compare CIBERSORT relative and absolute scores, we calculated the true amount of infiltration as two separate measures: “tissue” and “immune cell”. In the former, true amount of infiltration is calculated as the percent of reads from the immune cell type in the entire sample. In the latter, the true amount of infiltration is calculated as the percent of reads from the immune cell type in the immune content of the sample.

CIBERSORT was used to compute relative and absolute deconvolution scores of all synthetic mixes. All scores, regardless of generation process, were correlated with the true amount of infiltration in both the “tissue” and “immune cell” scenarios to quantitatively assess the differences.

### Merging cell subtype estimates into single scores

The CD4+ T cells category for CIBERSORT outputs reflects the sum of the “T cells CD4 naïve”, “T cells CD4 memory resting”, and “T cells CD4 memory activated” categories that are a part of the given LM22 reference matrix in CIBERSORT. The CIBERSORT “Macrophage” category represents the sum of the “Macrophages M0”, “Macrophages M1”, and “Macrophages M2” categories in the LM22 matrix. For the xCell analyses, CD4+ T cell scores were calculated by summing the scores from “CD4+ memory T-cells”, “CD4+ naive T-cells”, “CD4+ T-cells”, “CD4+ Tcm”, and “CD4+ Tem”. Macrophage scores were calculated using “Macrophages”, “Macrophages M1”, and “Macrophages M2”. Lastly, CD8+ T cell xCell scores were calculated by summing “CD8+ naive T-cells”, “CD8+ T-cells”, “CD8+ Tcm”, and “CD8+ Tem”.

### Visualizing cellular heterogeneity estimates across GTEx

Dendrograms representing the degree of similarity in immune composition across the 48 tissues in the GTEx dataset with at least 70 samples were generated for each of the 3 deconvolution methods. For each method, the median value of estimated immune content for each cell type was computed within each tissue. For the xCell deconvolution, 34 immune cell types were used, and for both Cibersort Absolute and Relative, 22 cell types were used. The heatmaps were drawn in R using *pheatmap*^61^, with Euclidean distance metric and with the “complete” linkage method. The xCell, Cibersort Absolute, and Cibersort Relative plots used maximal values of 0.05, 0.5, and 0.3, respectively.

We plot the mean cell type score across all tissues, separately for each deconvolution method, in heatmaps sorted by the mean CIBERSORT-Absolute scores for neutrophils, macrophages, CD4+ T cells, and CD8+ T cells (Supplementary Figure 1). We compute pairwise correlations between all tissue × cell type phenotypes (eg. compare CD4+ T cells in sun exposed skin tissue individually with neutrophils, macrophages, CD4+ T cells, and CD8+ T cells in each tissue). We plot these pairwise correlations using heatmaps (Supplementary Figure 2). We visualized a single cell type across all tissues for each deconvolution method using boxplots, sorted by the CIBERSORT-Absolute cell type score (Supplementary Figures 6-9).

Lastly, we used t-SNE to visualize immune content within a single tissue type and identify whether any clusters exist. We used scatterplots to visualize the two components and colored each point (which represents a unique sample/individual) by measured CD8+ T cell content.

### Filtering infiltration phenotypes for statistical analysis

To reduce the number of tests while focusing on informative phenotypes, we further limit our next analyses to cases where the cell type is abundant in the tissue/sample type and statistical methods could be reliably powered. We filter the tissue × cell type (infiltration) phenotypes to only those that have:

1. a sufficient sample size of N > 70 (matched genetic, expression, and covariate information) (similar to GTEx threshold)
2. consistent overall infiltration of immune cells in that tissue type (>50% of CIBERSORT relative deconvolutions have p < 0.50 (null hypothesis is that no immune cells from the reference are in the sample) (p = 0.50 observed previously^19^)
3. the specific immune cell type is a substantial part of the average immune content in that tissue (> 5% mean abundance in all CIBERSORT relative deconvolutions of the tissue) (>5% cutoff observed previously)^19^
4. CIBERSORT-Absolute and xCell scores do not disagree agree with each other (no significantly negative correlation)

While we were interested in studying regulatory T cell infiltration, this cell type would not pass the 3rd filter and so was removed from analysis. This leaves a total of 73 tissue × cell type combinations, which we refer to as our infiltration phenotypes.

### Analyzing principal components of gene expression profiles

Principal component analysis was performed on the processed gene expression matrix for each tissue separately. A linear regression analysis was fit between each infiltration phenotype and each of the first four principal components in that tissue (one-by-one). The p-values across all models were adjusted using Benjamini & Hochberg’s false discovery rate (*FDR*) correction^21^. We then identified the minimum adjusted p-value for each infiltration phenotype, from its individual comparisons with the first four principal components. We tested at *FDR* = 0.1.

### Differential expression analysis of extreme infiltration patterns

Consensus clustering of samples was performed using the BioConductor package ConsensusClusterPlus^62^, modified with the *fastcluster* R package to obtain considerable speed-up of results^63^, with 2,000 resampling cycles and k-means clustering with Euclidean distance. The most robust number of clusters was then selected for each tissue-cell pair. Within a given tissue-cell pair, clusters were assigned labels of “hot” or “cold” based on the mean estimate of sample scores for the cell type of interest. This procedure was applied independently to each of the xCell, Cibersort Absolute, and Cibersort Relative deconvolutions of the 73 tissue-cell pairs. Samples consistently identified as hot and cold across all 3 sets were taken as “consensus” hot and cold samples and considered for differential expression.

Differential gene expression was performed between the consensus hot and cold samples for each tissue-cell pair using limma-voom^64^. To address class imbalance between the number of hot and cold samples, we required that there be at least 6 hot and 6 cold samples in each tissue-cell pair before proceeding with differential expression, for statistical reasons described previously^65^. This left 51 tissue-cell pairs with sufficient number of samples.

Further, to account for covariate effects, we considered age (numeric; binned into 10-year categories), sex (binary), death classification (categorical; 0, 1, 2, 3, 4), autolysis score (numeric), and sample collection site (categorical). Covariates were included in the design matrix if there were a minimum of 3 hot and 3 cold samples in each level of that covariate. However, if there were a single level of a covariate that did not feature hot samples, we required that there be no more than 5 cold samples for that level in order for the covariate to be included in the design matrix (Supplementary Table 3).

43 of the 51 phenotypes featured differentially expressed genes at Benjamini-Hochberg adjusted *P* < 0.01 and log fold-change > 2.0, after adjustment for covariates and filtering of immune gene signatures used by the xCell and Cibersort deconvolution algorithms. Canonical pathways significantly enriched in the genes of interest were identified by Ingenuity Pathway Analysis.

### Multiple regression model for identifying age and sex associations

A multiple linear regression model accounting for age (numerical; discrete, binned into 10-year categories), sex (binary), death classification (categorical; 0, 1, 2, 3, 4), autolysis score (numerical), and sample collection site (categorical) covariates was fit for each phenotype to estimate age and sex effects (***β***). This was repeated for each of the deconvolution methods, and the *p*-values were combined using Empirical Brown’s method^26^. This method uses a covariance matrix to combine dependent p-values, allowing the incorporation of distinct analyses from each deconvolution method. As a final step, Benjamini & Hochberg’s false discovery rate (*FDR*) correction^21^ was applied to adjust all age-covariate p-values, then to separately adjust all sex-covariate p-values.

### Merging dependent p-values from three deconvolution score analyses

The p-values from an analysis using CIBERSORT-Relative, CIBERSORT-Absolute, and xCell are all different (owing to separate scores) yet correlated (due to different methods at quantifying the measure). To calculate a single measure of significance from all three analyses, we used Empirical Brown’s method^26^. We calculate covariance matrices for each infiltration phenotype (eg. CD4+ T cells in sun exposed skin tissue, using both the CIBERSORT and the xCell scores) and employ Empirical Brown’s method to convert the three p-values into a single p-value. This framework allows incorporation of several different cell type estimation methods to capture unique infiltration patterns, before merging the results into single measures. The Empirical Brown’s method p-values are reported, but age and sex testing p-values were corrected using Benjamini & Hochberg’s false discovery rate (*FDR*) correction prior to assessing significance at ***α*** = 0.05.

### Dimensionality reduction of cellular heterogeneity in breast tissues

To visualize differences in breast tissue heterogeneity, t-distributed stochastic neighbor embedding (t-SNE)^66^ was applied to the full (original) 64-cell type infiltration matrix from xCell.

### Pre-GWAS: genotype and phenotype processing

Similar multiple regression models to the age/sex model were used for pre-analysis phenotype processing. This model contained identical covariates to those discussed previously, but also including the first three genotype-based principal components to control for any population stratification. Genotype-based principal component analysis was performed using the *--pca* function in *plink*^67^. Gene expression-based latent factors, such as PEER factors^68^, have been demonstrated to be a powerful approach to correct for unwanted noise and technical variation. However, as described previously, our gene expression-based principal components correlated strongly with deconvolution estimates. As a result, our gene expression-based principal components, which could drastically reduce statistical power and inflate false positive rates, were not included in the model. The model was used to calculate residuals, which were transformed into *z*-scores using a rank-inverse normal transformation as implemented in the *GenABEL*^69^ package in *R*. Genotypes were filtered by minor allele frequency (< 0.05), missingness (> 0.1), and Hardy-Weinberg Equilibrium p-values (<10^−6^). A total of 5.6 million SNPs remained for analysis.

### Genetic analysis and hypothesis testing

We tested for associations between genome-wide variants and infiltration phenotypes using a simple linear regression model and the likelihood ratio test as implemented in *GEMMA*^70^. This was repeated for each deconvolution method, returning three p-values for each SNP’s relationship with each infiltration phenotype (cell type score in a tissue). The three p-values were merged with the Empirical Brown’s method framework utilized previously in the age and sex testing. All SNPs below a genome-wide threshold of 5.0 × 10^−8^ were considered significant. A separate study-wide significance threshold was determined by correcting for the number of infiltration phenotypes tested (73): P < 6.8 × 10^−10^. The relationship between raw gene expression and infiltration estimates were tested using a linear model.

### GeneMania

GeneMania combines multiple biological databases with a weighted “guilt-by-association” algorithm to add relevant genes to the query list and identify network edges ^46^. The first GeneMania network was constructed using only *STAM2* as the query gene, adding up to 20 additional genes and 10 functional attributes.

For the second and larger network, ieQTLs (loci associated with an infiltration phenotype that are also GTEx eQTLs in that tissue) (GWAS catalog threshold: p < 10^−5^) were used to form a list of ieGenes (the target genes of ieQTLs). The list of genes (recorded in Supplementary Table 16) were uploaded to the GeneMania software to construct a network of the input genes. In this analysis, 15 relevant functional attributes were used to supplement 30 genes to the original query of 85 genes. ieGenes with no shared edges with any other ieGenes were removed. To quantitatively assess the connectivity of each newly added gene to the network, GeneMania computes a score which was used to rank and identify the most interconnected genes.

### Testing for eQTL enrichment in iQTLs across phenotypes

iQTLs (P < 10^−5^; listed in Supplementary Table 17) were tested for over-or underrepresentation of tissue-specific eQTLs using two approaches. In the first approach, a 2×2 table is created by assessing whether each SNP is a GTEx eQTL and whether each SNP is an iQTL (relaxed threshold: p < 10^−5^). A chi-square test was performed to test whether tissue-specific eQTLs were distributed non-independently in the iQTL results for each phenotype.

In the second approach, for each phenotype, we generated the list of *N* iQTLs and match each of the *N* variants with a list of similar variants, as determined by minor allele frequency (within 1%, as calculated using --*freq* in *plink* from all GTEx individuals’ genetic data) and the same number of variants in linkage disequilibrium (LD) (r^2^ > 0.2, as calculated in the 1000 Genomes Phase I EUR genetic data^71^ and downloaded from Haploreg v4^30^). (We note that the threshold requiring the number of variants in LD to be identical is relaxed to plus-minus five variants-in-LD when no such variants exist.) We use these lists to generate 100 permutations. For each permutation, we randomly sampled 1 matched SNP for each of the *N* iQTLs. From the list of *N* randomly sampled SNPs, we calculated the proportion of SNPs that are tissue-specific eQTLs. From these 100 permutations, we calculated 100 eQTL proportion measurements. We then calculated the mean proportion, which we refer to as *q*. We let *x* be the # iQTLs that eQTLs in that tissue (tissue-specific eQTLs). Lastly, we performed a two-sided binomial test with *x* equal to the number of successes, *N* equal to the number of trials, and *q* equal to the hypothesized probability of success. We tested the null hypothesis that the observed ieQTL proportion is significantly different than random sampling. This approach is summarized in Supplementary Figure 17.

## Acknowledgements

We would like to thank members of the Clark laboratory and members of the Elemento laboratory for discussion surrounding this project and Zakieh Tayyebi for help designing an body-infiltration map (Figure 1a). We thank the GTEx donors for their contributions to science and the GTEx consortium for generating raw and analyzed data for researchers to access.

## Author contributions

A.R.M., A.G.C., and O.E. conceived and designed the study. A.R.M and M.U. performed the GTEx analysis. A.R.M., M.U., and A.V. designed and performed the simulations to test the deconvolution methods. A.V. and B.B. aided in developing the analysis methods and provided samples for the simulations. J.M. provided access to GTEx data. A.R.M., M.U., A.G.C., and O.E. wrote the manuscript. A.G.C. and O.E. supervised the study. All the authors reviewed and approved the manuscript.

