## Supplementary Information for "Age, Sex, and Genetics Influence the Abundance of Infiltrating Immune Cells in Human Tissues"

**Supplementary Notes:**

**Theoretical comparison of relative versus absolute scores.**

We motivate incorporating relative and absolute outputs from CIBERSORT into analyses by describing how both focus on capturing separate aspects of cellular heterogeneity.

Let $x$ be the true % of the sample that is immune cell type x.
Let $y$ be the true % of the sample that is immune cell type y.
Let $z$ be the total % of the sample represented by immune infiltration.

3 different samples with infiltration: $(x_1,y_1)$, $(x_2,y_2)$, $(x_3,y_3)$ = (10,0), (10,10), (20,0).
Therefore, $(z_1, z_2, z_3) = (10, 20, 20)$.

Let $\kappa$ be the relative percentage (%) of cell type $x$ to the infiltration profile. This can be described generally by the following expectation:

$$E[K] = x/z \tag{1}$$

CIBERSORT Relative measures the relative % of cell type $x$ to the other cell types in its reference (LM22, the infiltration landscape), e.g. $\kappa$. Using Equation 1, (E[$\kappa_1$], $E[\kappa_2]$, $E[\kappa_3]$) = (1, 0.5, 1). CIBERSORT Absolute scales this relative score to reflect the amount of overall infiltration. Let $\pi$ represent this scaling factor. For the purpose of this example:

$$E[\pi] = z/min(z_1, z_2, z_3) \tag{2}$$

such that $(\pi_1, \pi_2, \pi_3) = (1, 2, 2)$.
Let $\gamma$ be the CIBERSORT Absolute score, which should follow this expectation:

$$E[\gamma] = E[\pi]E[\kappa] \tag{3}$$

Using Equation 3:

$E[\gamma_1] = E[\pi_1]E[\kappa 1] = (1)(1) = 1$
$E[\gamma_2] = ... = (2)(0.5) = 1$
$E[\gamma_3] = ... = (2)(1) = 2$

Now, $E[\gamma_1] = E[\gamma_2] < E[\gamma_3]$ while $E[\kappa_2] < E[\kappa_1] = E[\kappa_3]$. Absolute and relative scores are quantifying different measures of the immune infiltration: immune cell relative composition (%) versus immune cell absolute counts.

Furthermore, CIBERSORT Relative calculates a better measure of the relative composition when $E[\pi_1] = E[\pi_2] = E[\pi_3]$. This is achieved by not including the scaling factor $\pi$:

$$Var[\gamma] = Var[\pi\kappa] > Var[\kappa] \tag{4}$$

Not including $\pi$ reduces the variance of the output, since $\pi$ has non-zero variance.

**Deconvoluting gene expression profiles from GTEx.**
We performed a simple merging of relevant cell subtypes to calculate larger group cell type scores (eg. in CIBERSORT, “Macrophages M0”, “Macrophages M1”, and “Macrophages M2” were summed into a single “Macrophages” score) (see **Methods**). We analyzed the reliability of cell type estimation between methods. We noted that mean cell scores for immune cell types of interest demonstrated marked heterogeneity between deconvolution methods, potentially capturing diverse infiltration patterns (Supplementary Figure 1). However, pairwise correlation between all infiltration phenotypes showed that major correlations are conserved between all methods, indicating that the methods still generally agree with each other (Supplementary Figure 2). Additionally, infiltration driven by the absolute amount of immune content is better captured in absolute outputs, as seen by more positive correlations across infiltration phenotypes in absolute scores compared to relative scores (Supplementary Figure 2). Finally, we were encouraged to use all three methods given the differences in capturing infiltration signals. For example, relative scores could be very similar but the absolute scores very different due to a higher amount of total infiltration (Supplementary Figure 3a). Individuals GTEX-14LLW and GTEX-17F96 had high proportions of neutrophil content in lung tissue according to CIBERSORT-Relative. However, a ~50% larger scaling factor for GTEX-17F96 resulted in different CIBERSORT-Absolute scores. xCell had even larger differences between the two individuals, with over twice as high estimated neutrophil content in GTEX-17F96. Similarly, xCell and CIBERSORT may also compute contrasting results for a single sample (Supplementary Figure 3b).

**Hierarchical clustering analysis of GTEx tissues.**
In our hierarchical clustering across the three cell type scoring methods, many brain tissues cluster together, as well as the tissue pairings of sun exposed and unexposed skin tissue and the tissue pairings of coronary and aorta arteries. For many other tissues, the nearest-neighbor pairing was not exact across the 3 deconvolution methods but demonstrated concordance through consistent clustering along the same branches, such as vagina and uterus tissues from the female urogenital tract or the heart tissues. In contrast, there were several tissues without consistent pairings in nearest-neighbor or overall branch, such as lung and liver, which reflect differences between the cell type estimation methods. We noticed different clustering among sigmoid and transverse colon tissue. Transverse colon had a much greater immune presence compared to sigmoid colon samples (scaling factor from CIBERSORT - Absolute: 1.96 in transverse, 1.08 in sigmoid colon samples), with greater detection of CD8+ T cell content (Supplementary Figure 5) (and insignificant correlation of CD8+ T cells between sigmoid and transverse tissue samples from the same individual).

**Analysis of inter-tissue differences across GTEx tissues.**
We observed the highest neutrophil scores in whole blood, spleen, and lung tissues across each deconvolution method (Supplementary Figure 8). While neutrophils dominate whole blood content, and therefore are expected in high frequencies within spleen tissue, neutrophils are frequently observed in lung tissue inflammation[1]. We also find the small intestine samples from the terminal ileum to have high CD8+ T cell content (Supplementary Figure 6), which makes

sense given the tissue's key role in digestion and encountering foreign microorganisms and possible infection. This also matches a gradient of increasing CD8+ T cell content from the rectum to further down the gastrointestinal tract (transverse and sigmoid colon), which corroborates previous clinical observations[2] and our own findings (Supplementary Figure 6).

**Analysis of differential expression between inflamed and non-inflamed samples**

Of the 10 CD8+ T-cell infiltration phenotypes, the most common DEGs in hot clusters are the CD8 lymphocyte marker genes *CD8A*, *CD8B*, and *CD3D*, the chemokine *CCL5*, immune receptor *KLRK1*, cytolytic marker *GZMK*, and immunomodulatory cell surface marker *SLAMF7* (all 9/10). With the exception of *SLAMF7*, each of these genes is used in the deconvolution algorithms' reference gene sets. The other most common DEGs not present in the reference sets include *THEMIS* and *TRBC2* (both 8/10, Supplementary Table 5), which are T cell-specific proteins, the former playing a critical role in CD4+ CD8+ thymocyte development, and the latter consisting of the constant region of the T cell receptor beta chain[3]. IPA further corroborated differential lymphocyte infiltration and activity, with the most frequent dysregulated pathways in hot clusters pertaining to leukocyte extravasation signaling (Supplementary Table 6), with the most commonly activated upstream regulator as IL1-beta (Supplementary Table 7). The gene ontology of most commonly activated pathways again pertained to leukocyte and lymphocyte chemotaxis (Supplementary Table 8).

For the 9 CD4+ T-cell infiltration phenotypes, the most DEGs most commonly expressed T-cell receptor-CD3 complex transcripts *LCK*, *UBASH3A*, *CD3E*, and *CD3G*; immunoglobulin genes *IGKV1-6*, *IGKV1-9*, *IGLV3-21*, and *IGHV1-24*; immune adhesion molecules *CD2* and *CD6*, and *THEMIS* (all 7/9, Supplementary Table 5). With the exception of *THEMIS* again, the genes are defined in the deconvolution algorithm reference sets for immune content estimation. The most frequently dysregulated pathways for CD4+ T cell included Th1 signaling, PKC-theta signaling in T lymphocytes, and *CD28* signaling in T Helper Cells (Supplementary Table 6), with *TNF* as the most commonly predicted activated upstream regulator (Supplementary Table 7). For CD4+ T cells, the diseases and functions ontology of activated pathways was associated with chemotaxis, cell migration, and cell homing (Supplementary Table 8).

Of the 27 most common DEGs across the 3 neutrophil-tissue pairs, 13 were present in the reference gene sets. The remaining 14 transcripts, 3 have been identified in neutrophil cytokinesis (*S100A12*, *S100P*, *PROK2*), 2 neutrophil degradative enzymes (*ARG1*, *LIPN*), 4 myeloid cell transmembrane receptors (*IL1R2*, *MS4A3*, *FCAR*, *CD177P1*), 2 molecules generally linked to immune activation (*ORM1*, *DUSP13*), and, 3 with uncharacterized function but known to be highly expressed in myeloid cells (*CTB-61M7.2*, *CTC-490G23.2*, *AC007278.2*). The most commonly dysregulated pathway was cAMP-mediated signaling (Supplementary Table 6), with activated upstream regulators being *TGM2* and *CEBPA* (Supplementary Table 7), and gene ontology associated with chemotaxis and cell movement of granulocytes and neutrophils (Supplementary Table 8).

**Sex associations with breast tissue.**

Temporal dependent changes (for example, the transition from lactating states to non-lactating states and menopause status) have been associated with an altered T cell response[4], and T cells have been associated with lobule localization (with higher densities of CD8+ T cells compared to CD4+ T cells)[5]. Furthermore, female breast may harbor a higher population of antigen-presenting cells to protect against potential infections compared to male breasts, which are less exposed to infection (eg. mastitis). To further assess the differences in the immune content of breast samples between males and females, we applied t-distributed stochastic neighbor embedding (t-SNE)[6] to the 22 immune cell scores from CIBERSORT-Absolute. The two t-SNE components displayed visual differences in clustering between males and females (Supplementary Figure 12).

**Potential rs77155650 effects on lung infiltration.**

The rs77155650 locus was also associated with CD8+ T cells, CD4+ T cells, and macrophages in lung tissue samples ($P < 0.05$; Supplementary Table 10), but not at genome-wide significance.

***CUX1* relationship with neutrophil infiltration in lungs.**

In mice, the DNA binding activity of the CUX-1 protein is triggered by the *F2RL1*-encoded $PAR_2$ protein[7], which has been linked to inflammation effects[8]. The DNA binding of the CUX-1 protein transcriptionally activated the expression of *IL1A*, *MMP10*, and *COX2* downstream[7]. *IL1A* plays a central role in the immune response and inflammation. *COX2* expression is often induced in inflamed tissue[9] and cancers (especially lung tumors)[10]. The MMP-10 protein encoded by the *MMP10* gene is a part of the matrix metalloproteinase family (MMP). While MMP-10 has been linked to lung cancer progression[11], lung inflammation[12], and lung disease[13], the related MMP-9 is secreted by neutrophils and has a key role in lung inflammation[14]. Additional evidence in mouse models suggests that *CUX1* plays a role in neutrophil infiltration and inflammatory mechanisms[15]. Furthermore, proteinases released by neutrophils at inflammation sites alter $PAR_2$[16] and CUX-1[17] functions within a possible feedback loop. Potentially, the rs77155650 polymorphism could interact with *F2RL1*-triggered CUX-1 DNA binding and the downstream activation of *IL1A, MMP10,* and *COX2* for initiating neutrophil inflammation processes.

**Potential rs116827016 effects on artery infiltration.**

In the other 7 infiltration phenotypes from arterial tissue (from aorta: CD8+ T cells, CD4+ T cells, and macrophages; from coronary: CD8+ T cells, CD4+ T cells, and macrophages; from tibial: CD4+ T cells and macrophages), rs116827016 has a median p-value of 0.019 and minimum p-value of $5.9 \times 10^{-3}$, further suggesting infiltration effects (Supplementary Table 11).

***KCTD10*'s potential role in arterial inflammation.**

We note that a paralog of *KCTD10* is *TNFAIP1*. *TNFAIP1* has been associated with IL-5 levels, with its regulation linked to inflammation-associated angiogenesis[18]. Similarly, previous functional studies of homozygous *KCTD10* knockout mice showed severe defects in angiogenesis[19]. It is plausible that *KCTD10* shares many functions with *TNFAIP1*, and that

*KCTD10* plays a significant role to immune response within the vascular system. The rs116827016 haplotype possibly alters *KCTD10* expression, and changes in *KCTD10* expression could result in a modified crosstalk and immune function between the two related systems.

**GeneMania network analysis of *STAM2* function.**

To learn about the surrounding genetic interaction network and related functional pathways of *STAM2*, we queried *STAM2* in GeneMania (Supplementary Figure 14). GeneMania combines multiple biological databases with a weighted “guilt-by-association” algorithm to add relevant genes to the query and construct a network[20]. The resulting network of 20 genes and 7 functional attributes included *IL2* and *IL2*-mediated signaling events as key functional attributes (a key regulator of white blood cell activity). 19 of the 20 genes were linked to gene ontology (GO) functional annotations related to the EGFR and ERBB pathways. Epidermal growth factors drive cancer proliferation[21,22], and anti-EGFR inhibitors are associated with the immune response[23,24]. There were many other enriched GO functions, including toll-like receptor signaling, cellular response to peptides, and immune-response regulation by cell surface receptors.

**Phenotypes more enriched for eQTL overrepresentation rather than underrepresentation.**

eQTL-enrichment of results was more significant than eQTL-underrepresentation: the grouped -log10 p-values of eQTL-enriched phenotypes versus eQTL-depleted were significantly different for both methods (Mann-Whitney U test, $p = 0.03$ and $p = 2.8 \times 10^{-4}$ for Test 1 and Test 2 respectively).

**In-depth study of ieGenes using GeneMania.**

To attempt to draw functional conclusions from our genetic results, we used the gene expression associations from our ieQTLs. We constructed a GeneMania network by forming a list of ieGenes (genes whose expression is associated with the variant, as determined from GTEx analysis[25]) from ieQTLs with our relaxed iQTL threshold $p < 10^{-5}$ (GWAS catalog cut-off), developing a query of 85 genes. GeneMania supplemented 15 functional attributes and 30 additional genes to the original network, for a total of 115 genes (Supplementary Figure 16). We found that this network was enriched for pyrimidine biosynthesis and DNA repair functions (Supplementary Table 12). These functions help maintain healthy DNA, which is crucial for normal function and cancer avoidance. DNA damage sensors are tightly coupled with innate immune signaling mechanisms, since affected DNA repair may increase the rate of DNA damage, stimulating an immune response[26]. Furthermore, proliferation of immune cells is dependent on de novo nucleotide biosynthesis, which is a target of several existing immunosuppressive drugs[27]. To quantitatively assess the connectivity of each added gene to the network, GeneMania computes a score. The top seven added genes were (by rank): (1) *CAD*, (2) *DCTD*, (3) *TYMS*, (4) *RB1*, (5) *RBL1*, (6) *CREBBP*, and (7) *EP300*. *CAD, DCTD,* and *TYMS* are three key genes involved in pyrimidine biosynthesis and metabolism. Previous studies have linked *TYMS* to cancer growth[28,29], and inhibitors of its protein (thymidylate synthase) are widely used chemotherapy targets. The last 4 are involved in transcriptional mechanisms and epigenetic remodeling, with each linked to immune-related disorders. For example, *EP300* has been associated with lymphocyte percentages[30] and Crohn's disease[31].

**Supplementary Figure 1:** Heatmaps display the mean cell score of the given cell type (y-axis) in each sample type (x-axis) for three different deconvolution outputs (the relative estimates using CIBERSORT, the absolute estimates using CIBERSORT, and the estimates using xCell). Each column reflects the same sample type, which has been sorted by the sum of the CIBERSORT absolute scores for four cell types of interest.

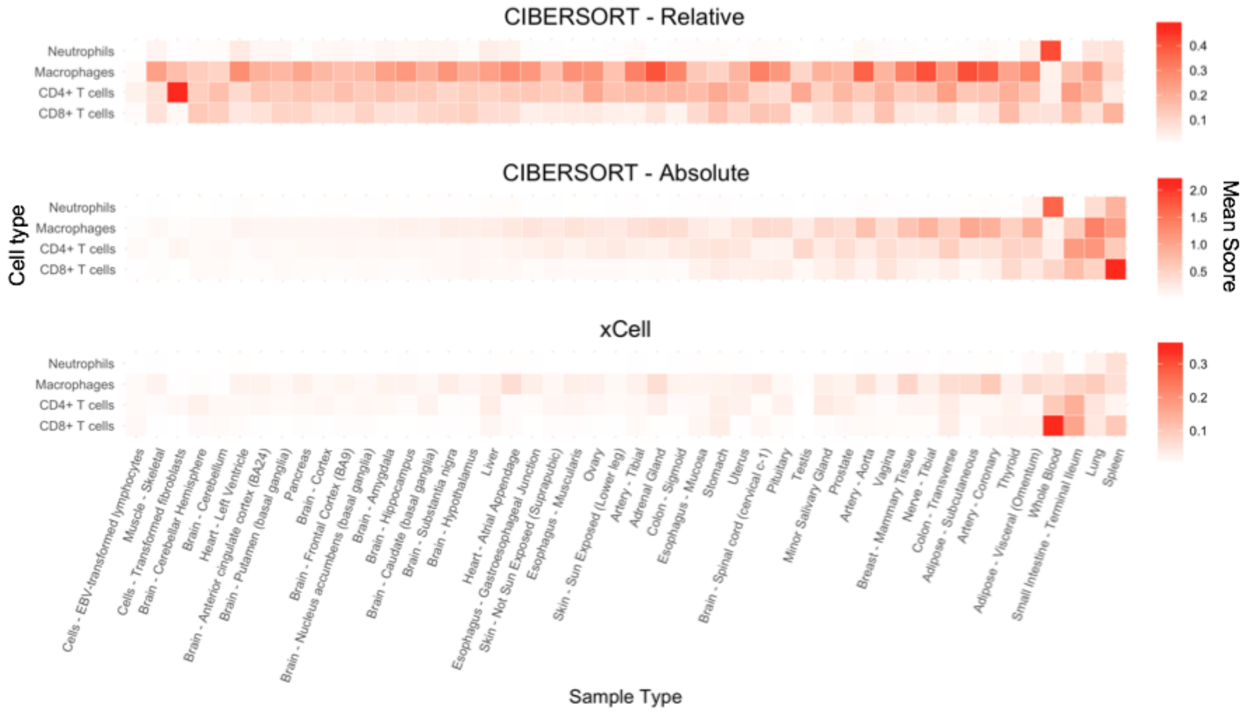

**Supplementary Figure 2:** Pairwise correlations were measured between all original infiltration phenotypes, spanning 4 cell types in each of 53 tissues. Each row and column is the same infiltration phenotype in each of the three plots. Correlations are displayed via heatmap, which show generally conserved correlations between methods. Additionally, absolute scores (CIBERSORT Absolute and xCell) show more positive correlations (redness) compared to relative scores (CIBERSORT Relative).

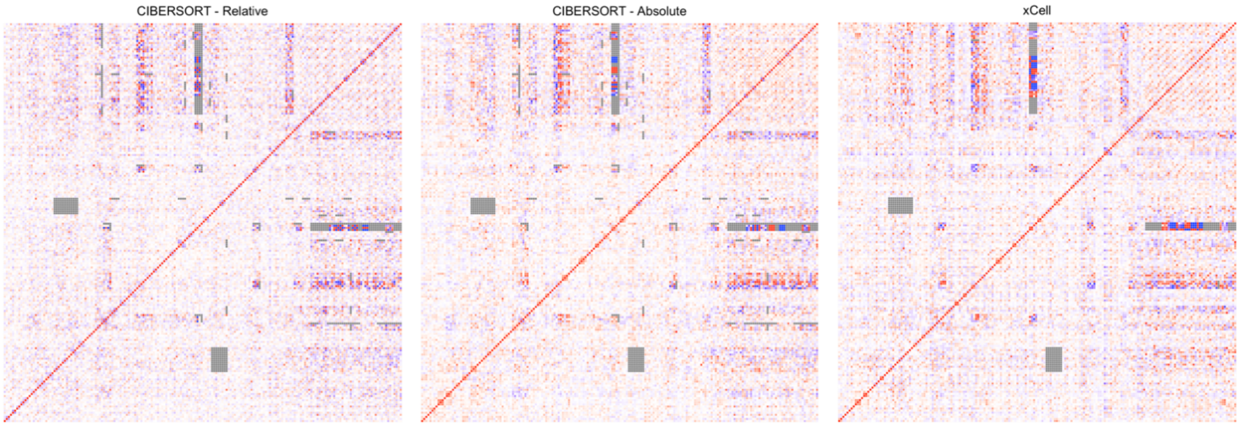

**Supplementary Figure 3:** Neutrophil content in lung tissue as estimated by CIBERSORT-Relative, CIBERSORT-Absolute, and xCell algorithms. (a) Individuals GTEX-14LLW (blue) and GTEX-17F96 (orange) have similar relative scores but different absolute scores. (b) Individuals GTEX-PLZ4 (blue) and GTEX-ZF2S (orange) have similar xCell scores, but very different CIBERSORT scores.

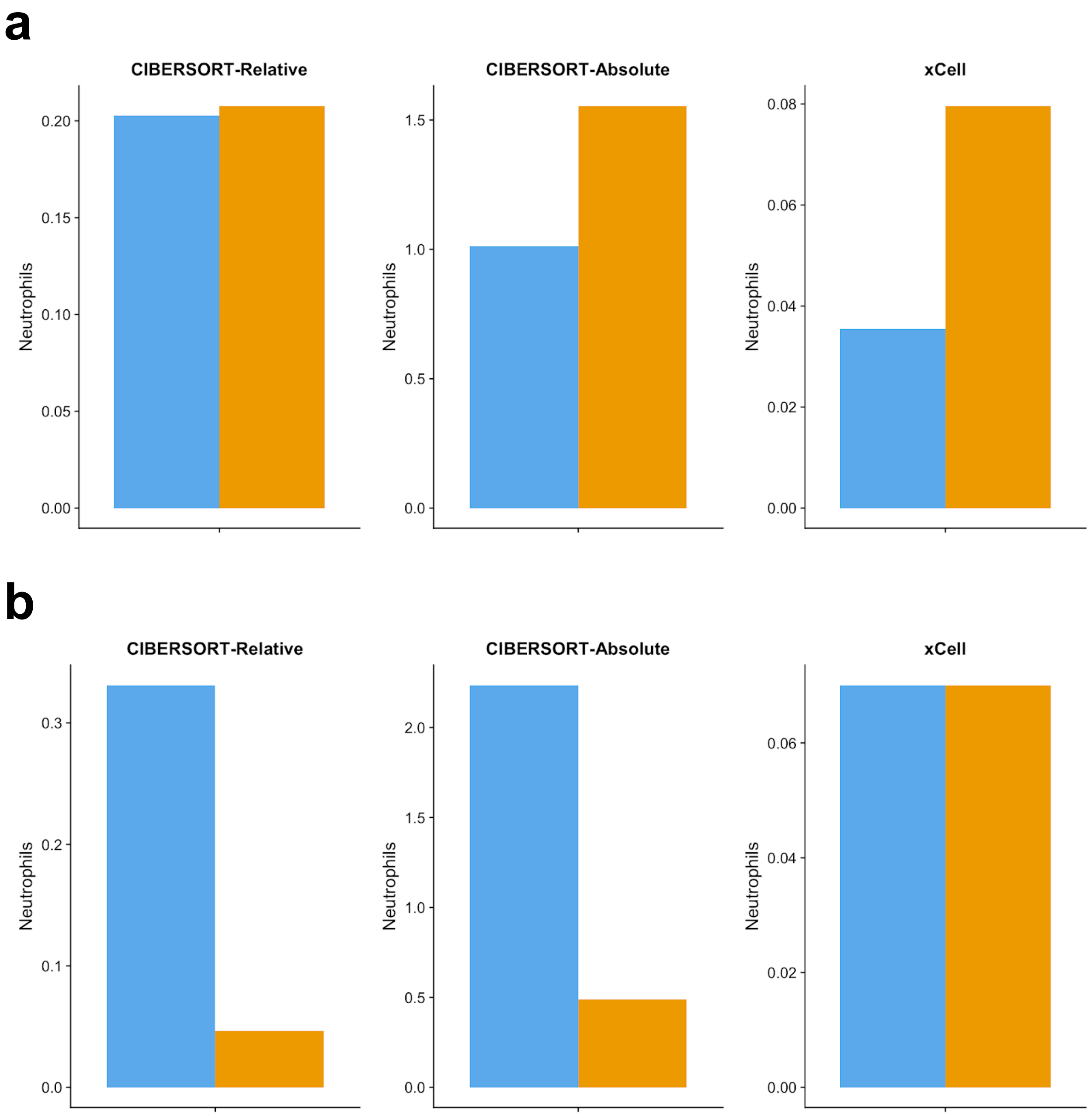

**Supplementary Figure 4:** Hierarchical clustering of GTEx tissues according to immune content (estimated by CIBERSORT-Relative). Heatmap displays cell type median scores, with upper bound of 0.5.

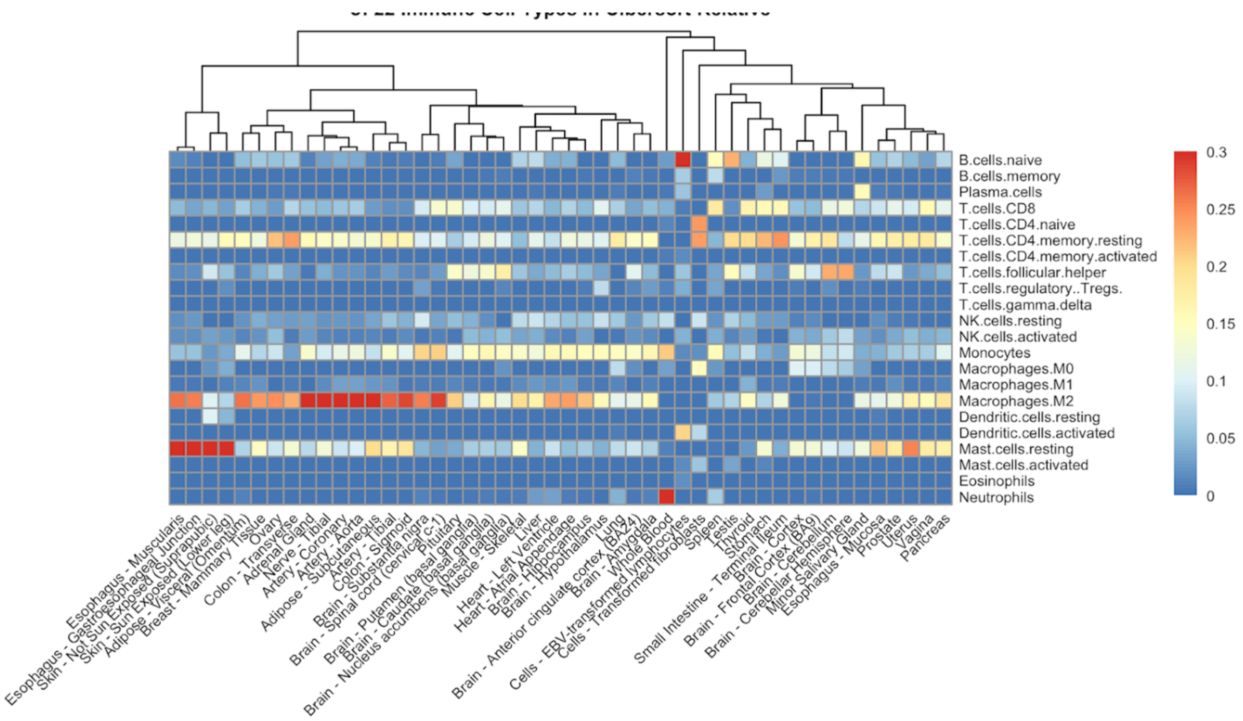

**Supplementary Figure 5:** Hierarchical clustering of GTEx tissues according to immune content (estimated by xCell). Heatmap displays cell type median scores, with upper bound of 0.05.

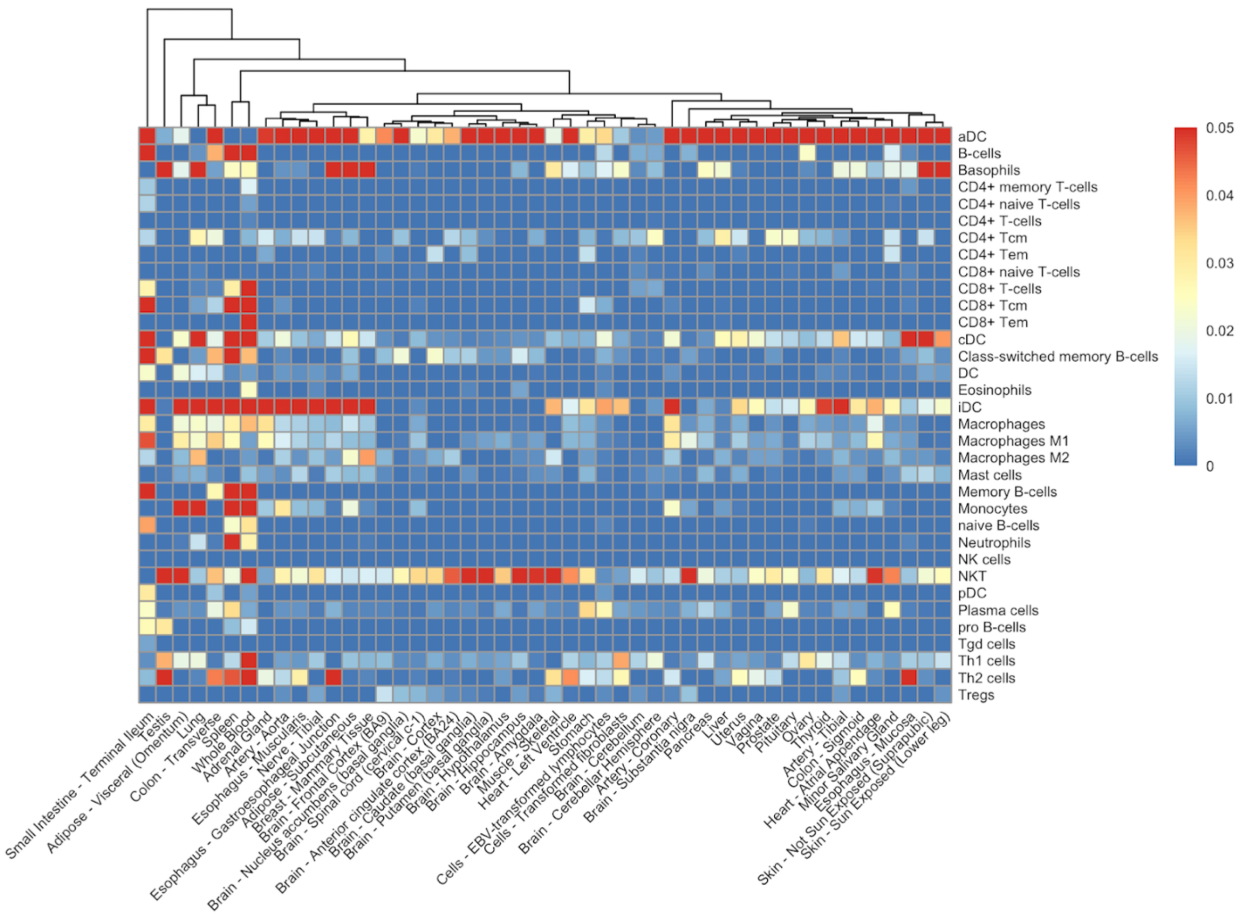

**Supplementary Figure 6:** Estimates of CD8+ T cell content in GTEx tissues, segmented by each deconvolution method. (a) is CIBERSORT-Relative, (b) is CIBERSORT-Absolute, and (c) is xCell. Tissues are sorted by median CIBERSORT - Absolute score.

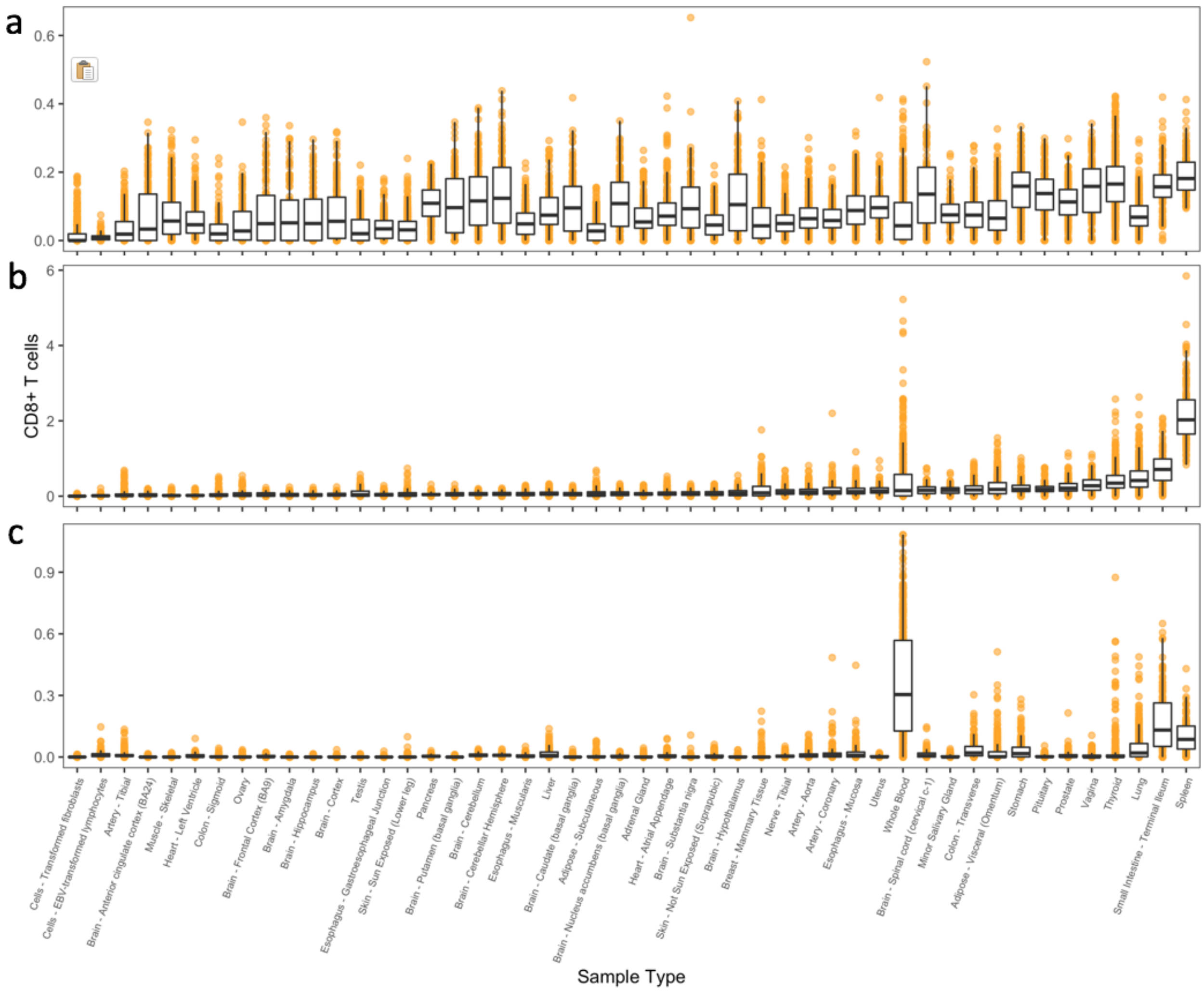

**Supplementary Figure 7:** Estimates of macrophages in GTEx tissues, segmented by each deconvolution method. (a) is CIBERSORT-Relative, (b) is CIBERSORT-Absolute, and (c) is xCell. Tissues are sorted by median CIBERSORT - Absolute score.

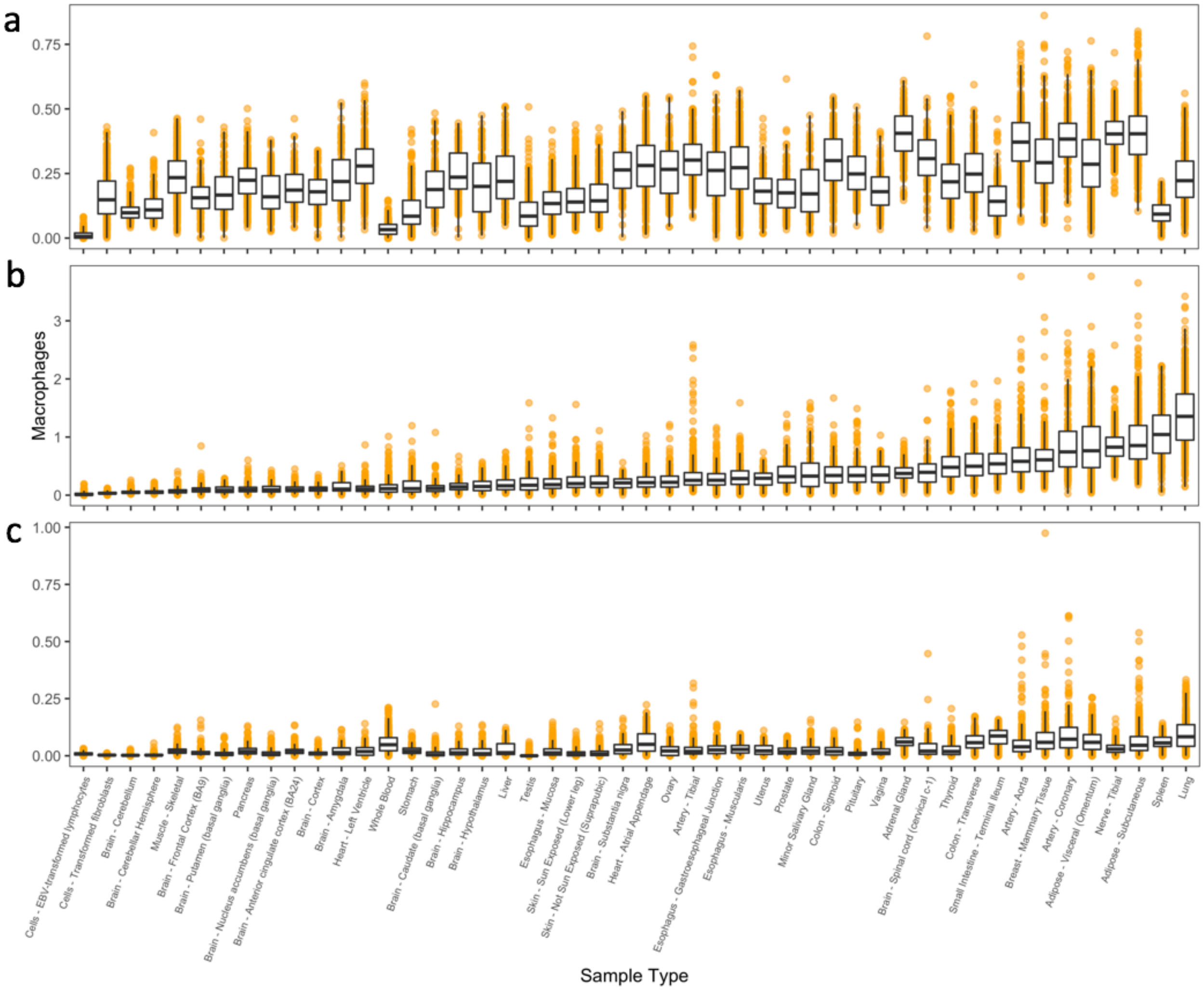

**Supplementary Figure 8:** Estimates of neutrophils in GTEx tissues, segmented by each deconvolution method. (a) is CIBERSORT-Relative, (b) is CIBERSORT-Absolute, and (c) is xCell. Tissues are sorted by median CIBERSORT - Absolute score.

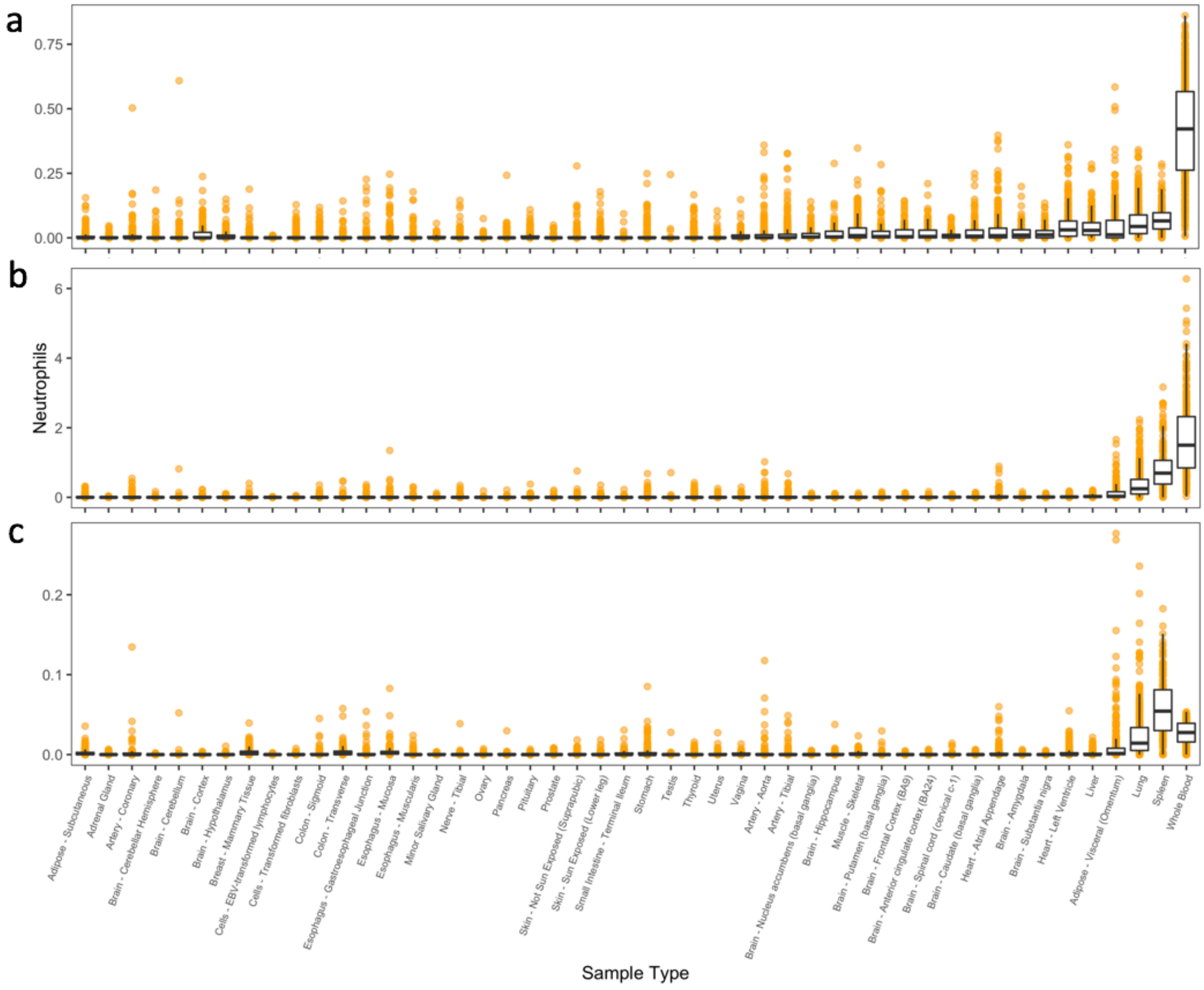

**Supplementary Figure 9:** Estimates of CD4+ in GTEx tissues, segmented by each deconvolution method. (a) is CIBERSORT-Relative, (b) is CIBERSORT-Absolute, and (c) is xCell. Tissues are sorted by median CIBERSORT - Absolute score.

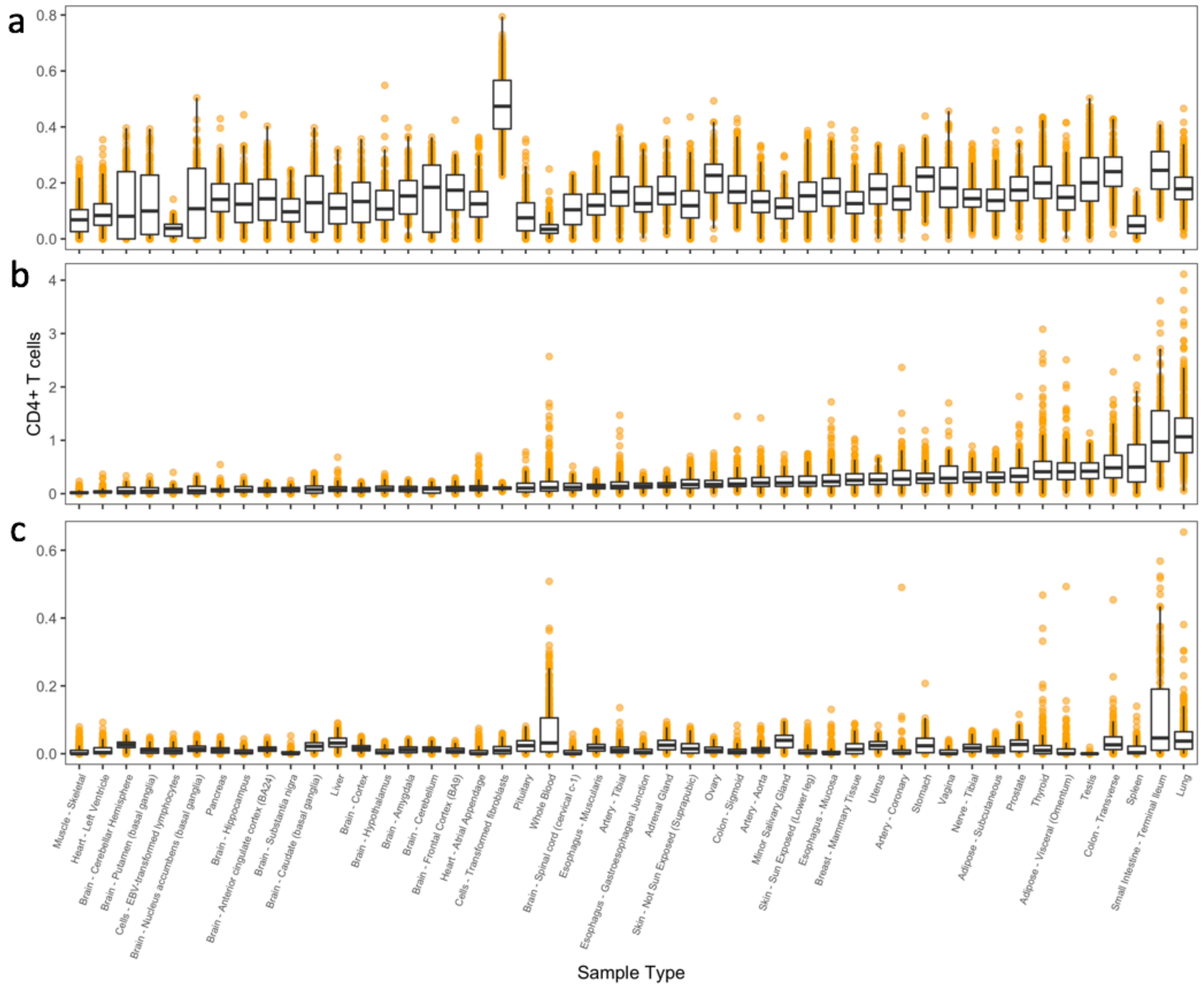

**Supplementary Figure 10:** t-SNE clustering of immune content within a single tissue type. t-SNE was performed on the 22 immune cell type scores from CIBERSORT-Absolute deconvolution. Each point represents a unique sample from a different individual, which has been colored by the amount of measured CD8+ T cell content (low content: blue; high content: red).

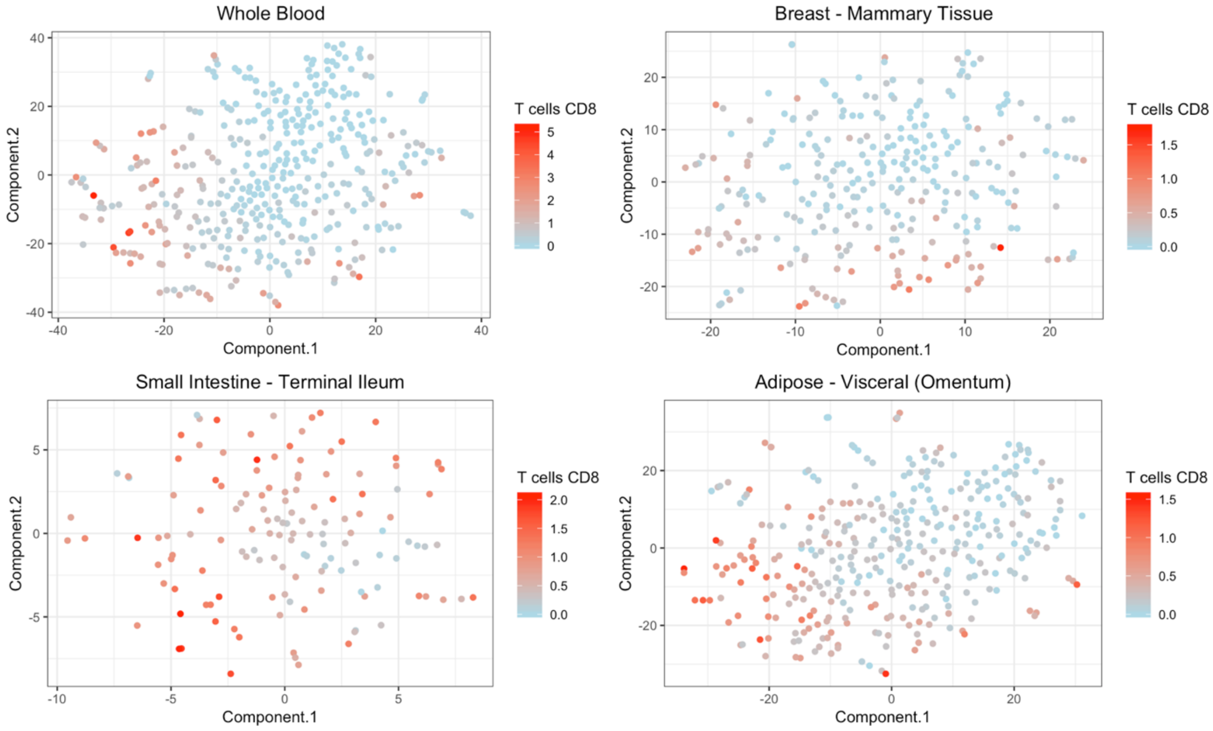

**Supplementary Figure 11:** Number of tissues where each individual is in the "hot" cluster, separated by cell type. Each bar plot is subset to individuals who are in at least one "hot" cluster of that cell type.

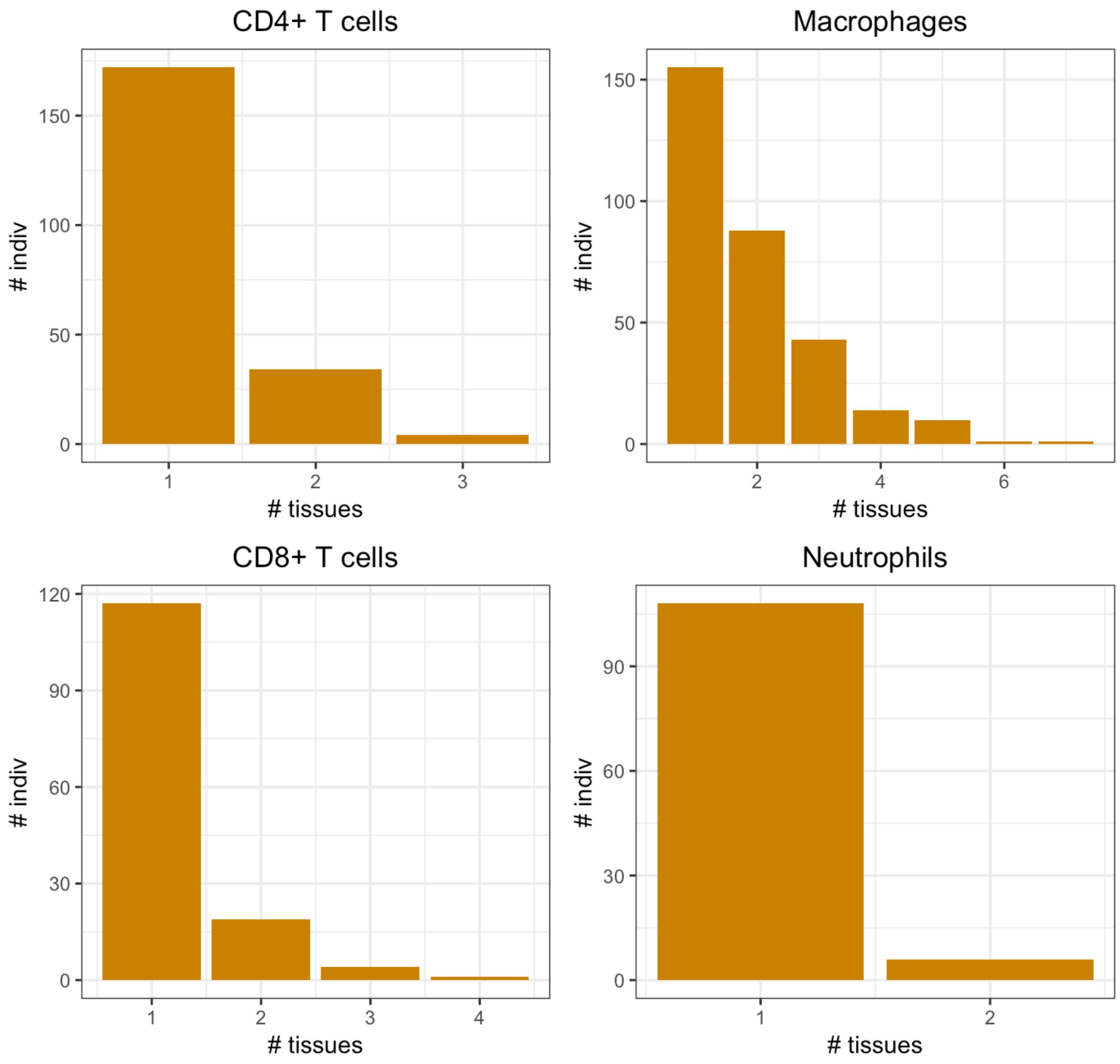

**Supplementary Figure 12:** t-SNE plot of immune content in breast tissue, calculated on the 22 immune cell type matrix from CIBERSORT - Absolute. Each point is a separate individual, colored by sex.

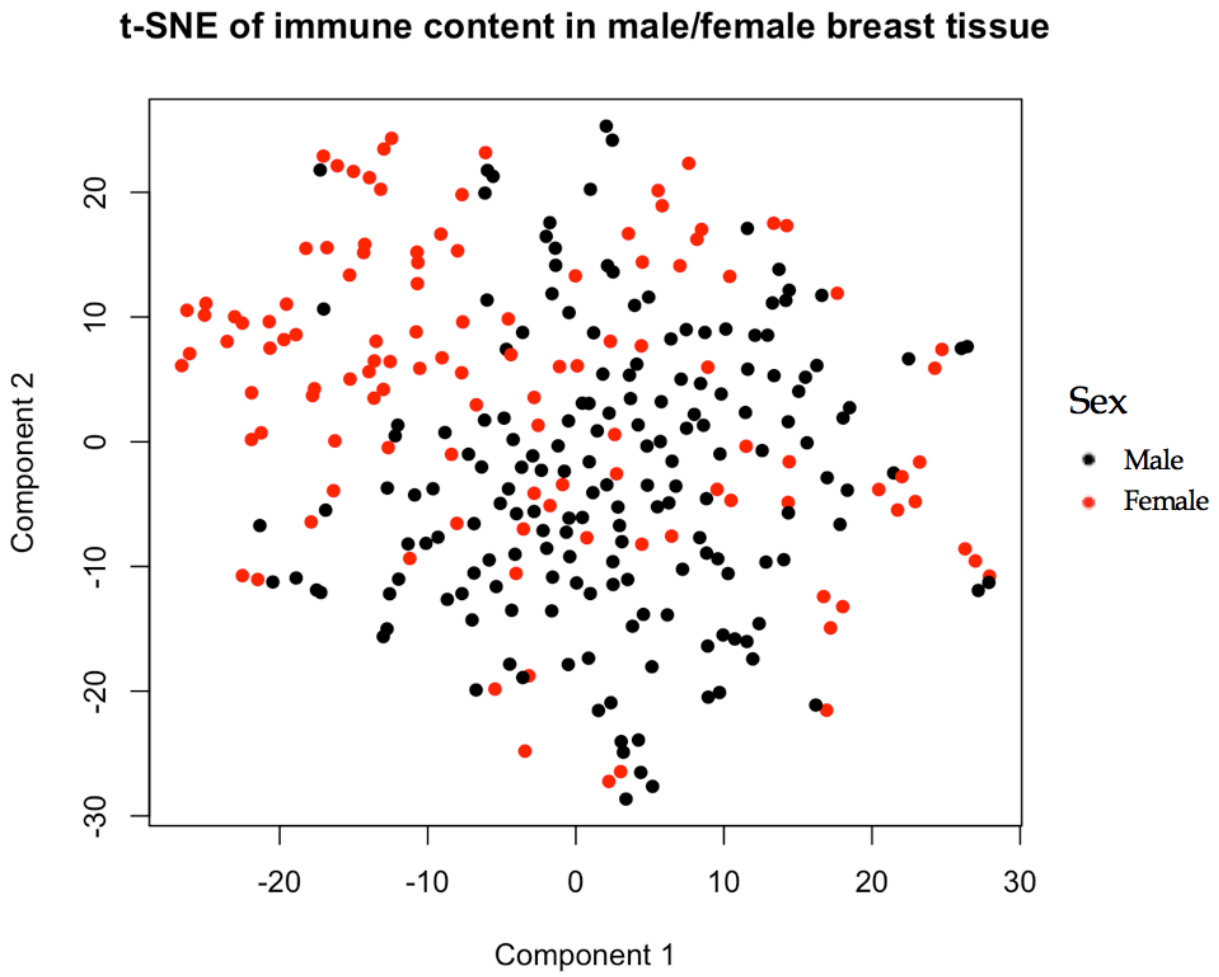

**Supplementary Figure 13:** KCTD10 gene expression across GTEx tissues. Expression is highest in tibial artery tissue.

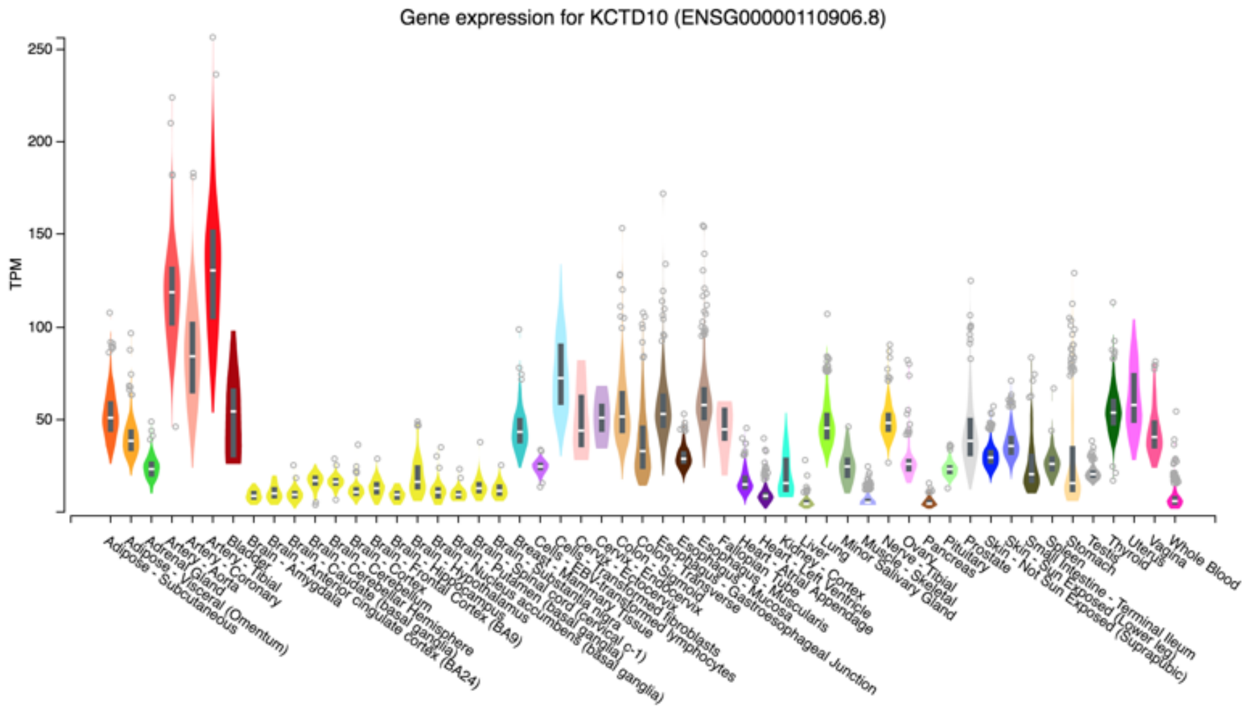

**Supplementary Figure 14:** STAM2 query in GeneMania, adding up to 20 genes and 10 functional attributes. Final network includes 20 total genes (19 added) and 7 functional attributes.

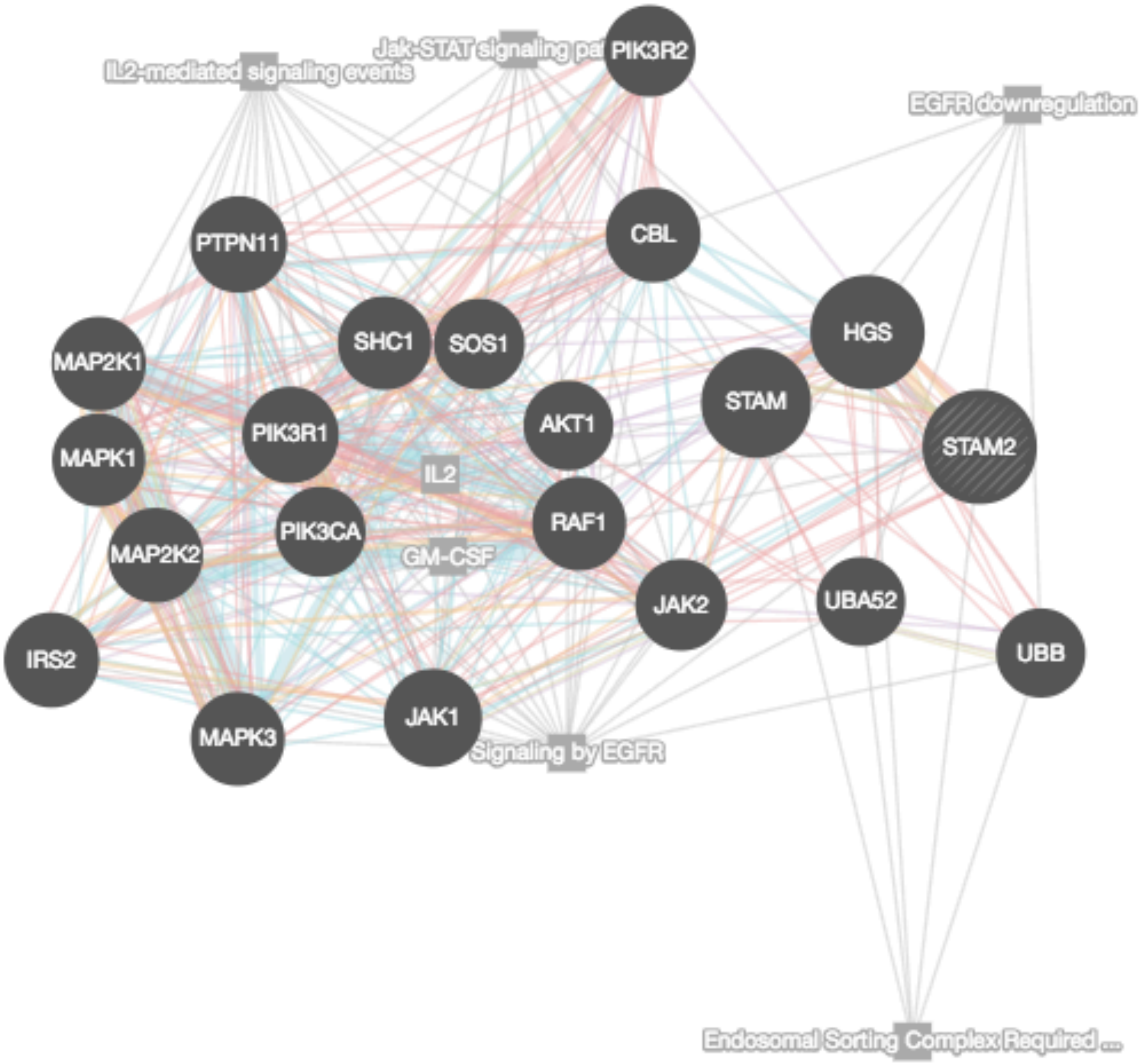

**Supplementary Figure 15**: eQTL enrichment in iQTLs across tissues, using test 2. Y-axis is -log10 p-values from binomial test and x-axis is each infiltration phenotype analyzed and sorted by significance.

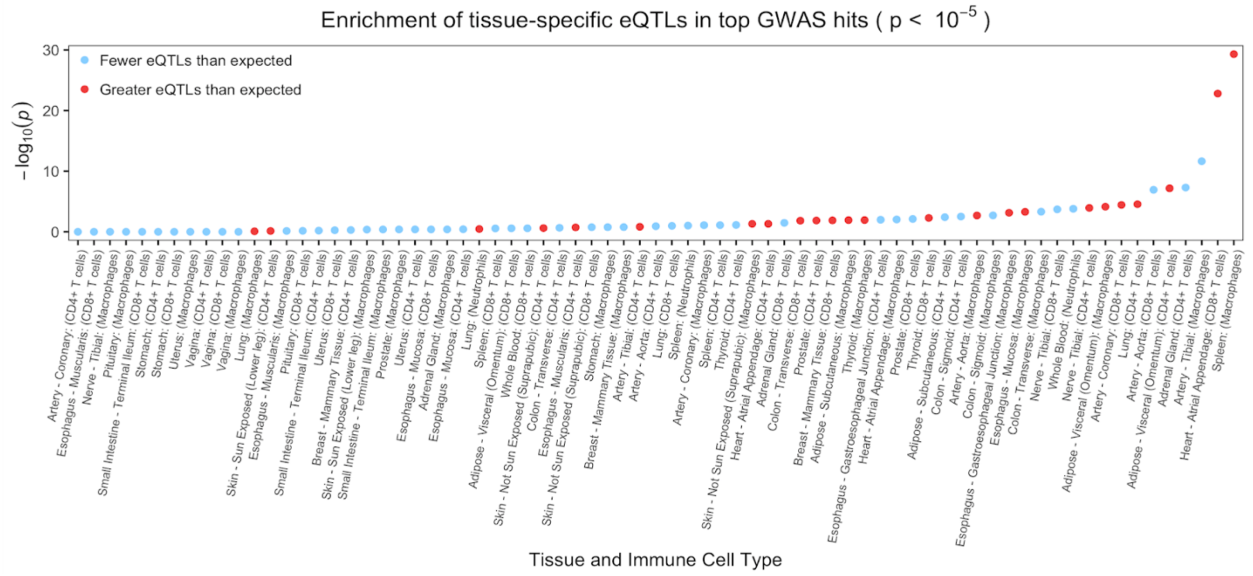

**Supplementary Figure 16:** GeneMania network of all ieGenes from ieQTLs that are associated with infiltration phenotypes at a relaxed $p < 10^{-5}$ threshold.

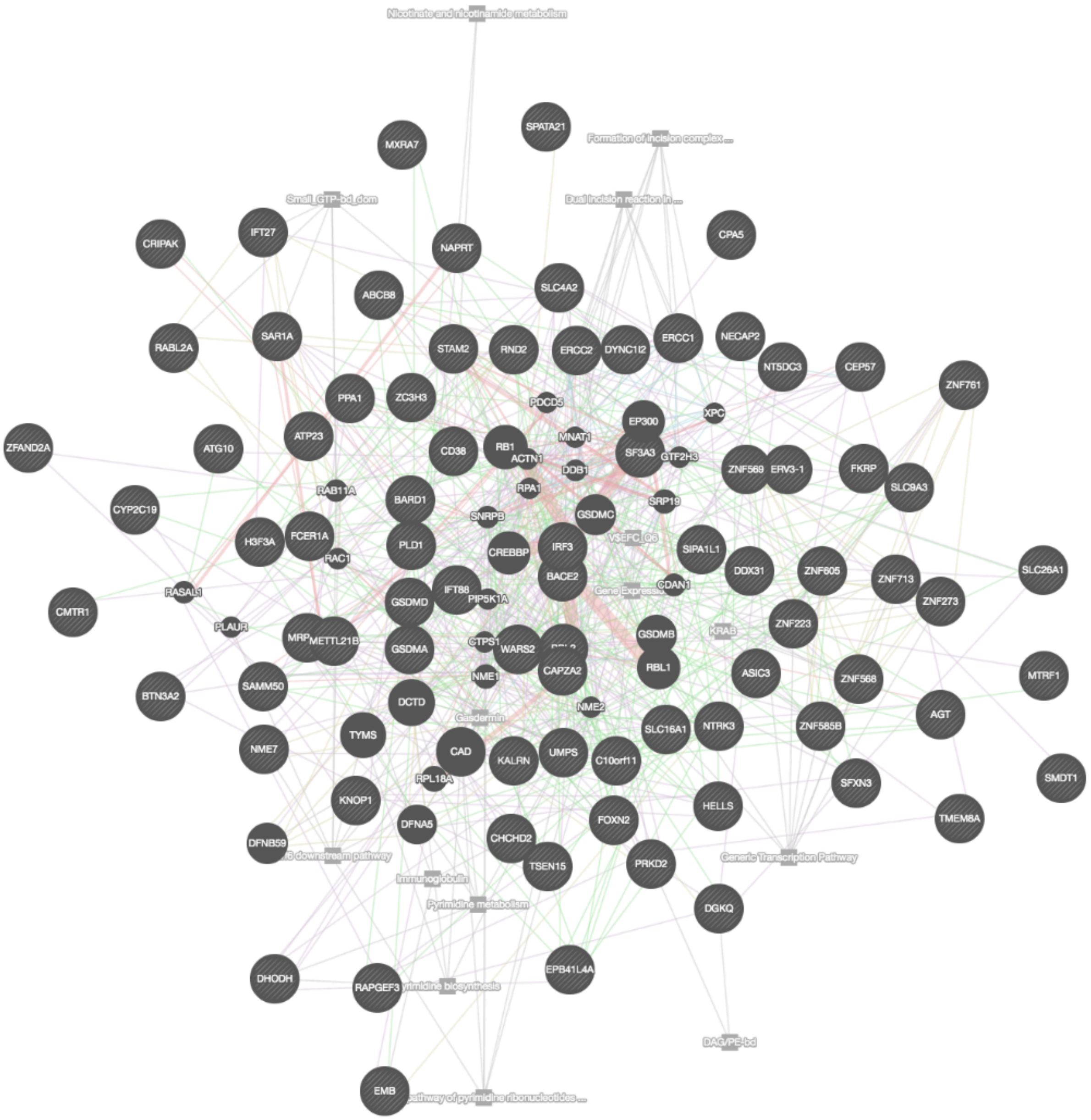

**Supplementary Figure 17:** Bi-partite networks of target ieGenes. Orange points are genes, yellow points are cell types. Edges represent significant association between ieQTL and cell type. Nodes shared between neighborhoods represent ieGenes that have ieQTLs associated with multiple cell types.

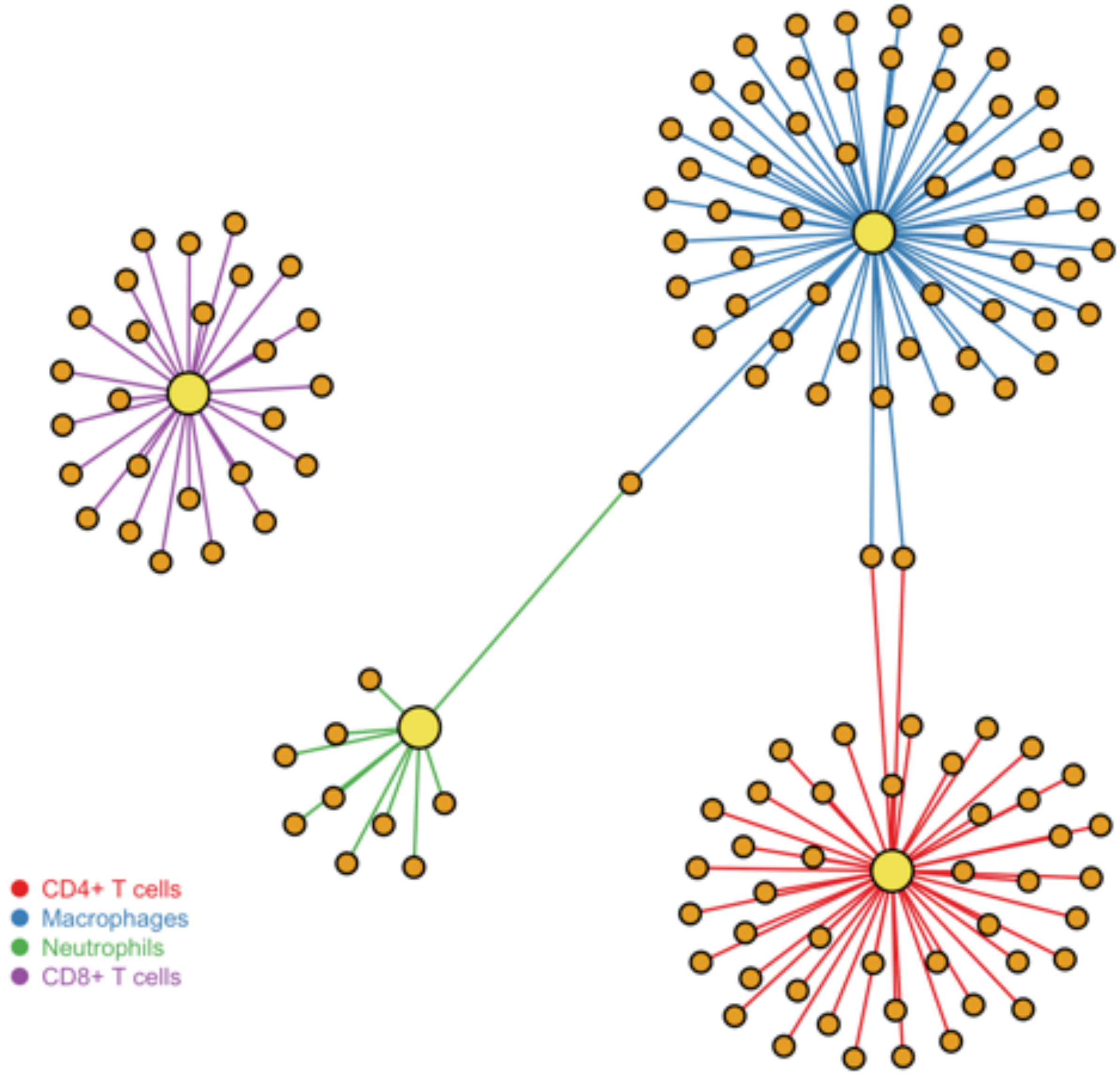

**Supplementary Figure 18:** Overview of test 2 for eQTL enrichment, based on a binomial test.

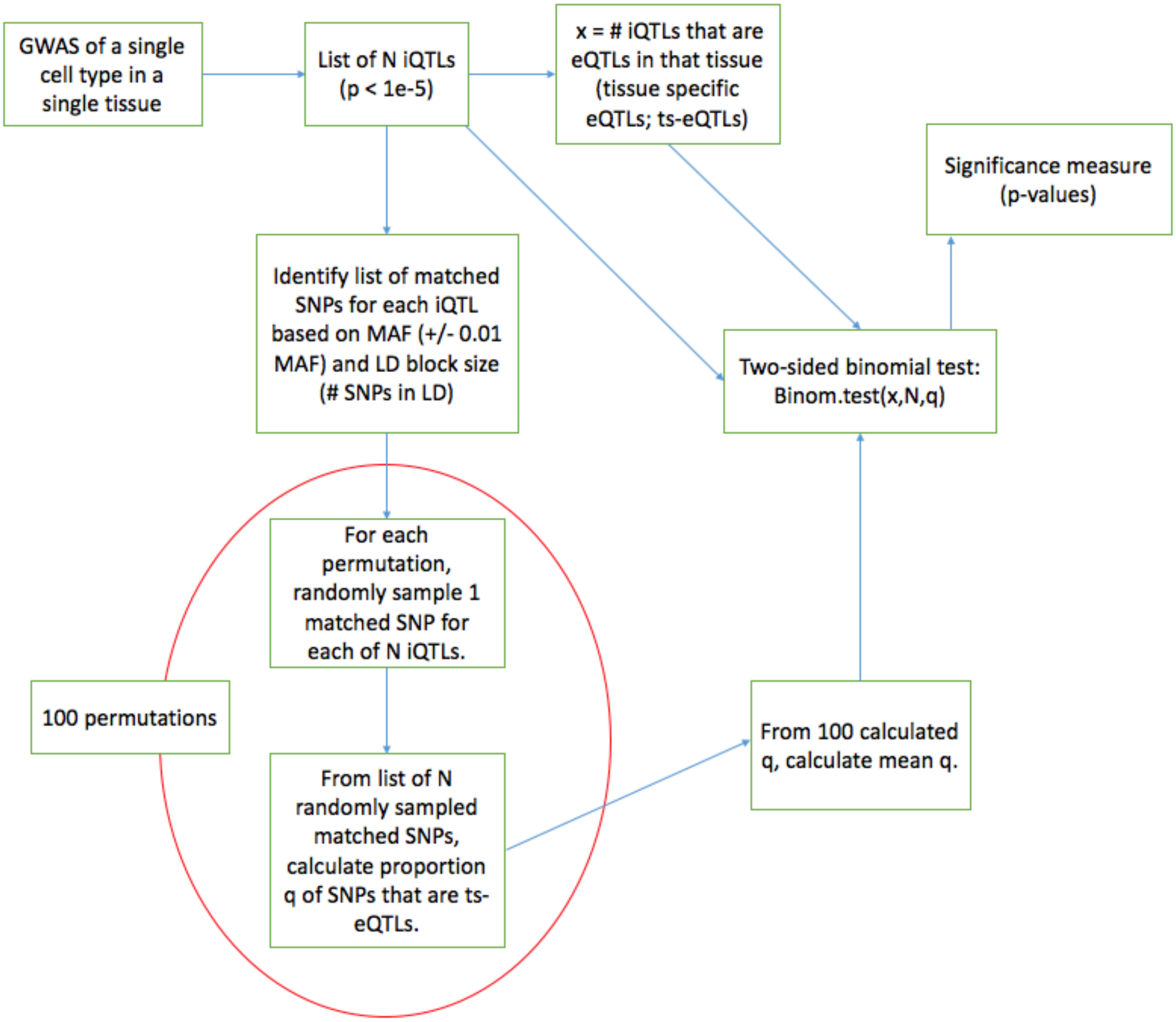

| Cell Type | Scenario | CIBERSORT Mode | Correlation |
|---|---|---|---|
| CD4+ T cells | Tissue | Relative | 0.7968766 |
| CD4+ T cells | Tissue | Absolute | 0.8884554 |
| CD4+ T cells | Immune Cell | Relative | 0.8389506 |
| CD4+ T cells | Immune Cell | Absolute | 0.8016166 |
| CD8+ T cells | Tissue | Relative | 0.6470465 |
| CD8+ T cells | Tissue | Absolute | 0.7115709 |
| CD8+ T cells | Immune Cell | Relative | 0.848441 |
| CD8+ T cells | Immune Cell | Absolute | 0.8001204 |

**Supplementary Table 1:** Correlation of estimated scores (from CIBERSORT) versus true quantity of cell type. True quantity of cell type measured as % of whole sample (absolute infiltration) and % of immune content (relative proportionality).

**Supplementary Table 2:** Correlation of xCell scores with true amounts, as tested on the simulated synthetic mixes. xCell mode “Tissue-by-tissue” describes estimation on each tissue separately, while mode “Simultaenous” describes estimation on the full gene expression matrix (including different tissues at once). Default scores are not normalized (do not sum to 1). Last 2 rows represent performance of xCell in the “immune cell” scenario when the xCell scores were normalized.

| **Cell Type** | **Scenario** | **xCell mode** | **Correlation** |
|---|---|---|---|
| CD4+ T cells | Tissue | Tissue-by-tissue | 0.9581576 |
| CD4+ T cells | Immune Cell | Tissue-by-tissue | 0.6511046 |
| CD8+ T cells | Tissue | Tissue-by-tissue | 0.9037872 |
| CD8+ T cells | Immune Cell | Tissue-by-tissue | 0.449362 |
| CD4+ T cells | Tissue | Simultaneous | 0.9214696 |
| CD4+ T cells | Immune Cell | Simultaneous | 0.6087092 |
| CD8+ T cells | Tissue | Simultaneous | 0.8419683 |
| CD8+ T cells | Immune Cell | Simultaneous | 0.4063541 |
| CD4+ T cells | Immune Cell | Tissue-by-tissue | 0.7290438 |
| CD8+ T cells | Immune Cell | Tissue-by-tissue | 0.5955122 |

**Supplementary Table 3:** Assignment of samples from 51 infiltration phenotypes to hot and cold phenotypes, and covariate data used in differential expression. Each sample is labeled “hot” or “cold” based on consistent classification across the 3 deconvolution methods.

https://drive.google.com/drive/folders/19KvgtIgKzmTTUY5WSB3kP8ubUVtvK08T?usp=sharing

**Supplementary Table 4:** Covariates included in design matrix for differential expression for 51 infiltration phenotypes, with “1” entailing inclusion of covariate, and “0” corresponding to exclusion. Covariate inclusion was based on sufficient representation of each level of the covariate in hot and cold clusters.

https://drive.google.com/drive/folders/19KvgtIgKzmTTUY5WSB3kP8ubUVtvK08T?usp=sharing

**Supplementary Table 5:** Differential gene expression results from 43 infiltration phenotypes expressing at least 5 transcripts at cutoffs of log fold change > 2 and adjusted p value < 0.01.

https://drive.google.com/drive/folders/19KvgtIgKzmTTUY5WSB3kP8ubUVtvK08T?usp=sharing

**Supplementary Table 6:** IPA canonical pathway analysis for hot and cold clusters across 43 infiltration phenotypes.

https://drive.google.com/drive/folders/19KvgtIgKzmTTUY5WSB3kP8ubUVtvK08T?usp=sharing

**Supplementary Table 7:** IPA upstream regulator analysis for the hot and cold clusters across 43 infiltration phenotypes.

https://drive.google.com/drive/folders/19KvgtIgKzmTTUY5WSB3kP8ubUVtvK08T?usp=sharing

**Supplementary Table 8:** IPA gene ontology analysis for the hot and cold clusters across 43 infiltration phenotypes. Annotations of disease or functional association and corresponding dysregulated molecules are represented.

https://drive.google.com/drive/folders/19KvgtIgKzmTTUY5WSB3kP8ubUVtvK08T?usp=sharing

**Supplementary Table 9:** The 73 post-filtered infiltration phenotypes from CIBERSORT-Absolute were tested for association with the first four gene expression-based principal components using a linear model. All p-values (73 phenotypes * 4 PCs = 292 total tests) were adjusted using a FDR correction. The minimum p-value for each infiltration (4 total p-values) is displayed, along with the adjusted value.

*See SupplementaryTable9.txt.*
https://drive.google.com/drive/folders/19KvgtIgKzmTTUY5WSB3kP8ubUVtvK08T?usp=sharing

**Supplementary Table 10:**
Association of rs77155650 with immune cell quantities in lung samples. Raw p-values shown.

| **Cell Type** | **P-value** |
|---|---|
| CD8+ T cells | $1.0 \times 10^{-2}$ |
| CD4+ T cells | $1.4 \times 10^{-3}$ |
| Neutrophils | $9.7 \times 10^{-11}$ |
| Macrophages | $1.2 \times 10^{-2}$ |

**Supplementary Table 11:**
Association of rs116827016 with immune cell quantities from artery tissues.

| **Artery Type** | **Cell Type** | **P-value** |
|---|---|---|
| Aorta | CD8+ T cells | 0.40 |
| Aorta | CD4+ T cells | 0.11 |
| Aorta | Macrophages | $6.5 \times 10^{-3}$ |
| Coronary | CD8+ T cells | $1.9 \times 10^{-2}$ |
| Coronary | CD4+ T cells | $5.9 \times 10^{-3}$ |
| Coronary | Macrophages | $1.6 \times 10^{-2}$ |
| Tibial | CD4+ T cells | $6.6 \times 10^{-2}$ |
| Tibial | Macrophages | $3.9 \times 10^{-10}$ |

**Supplementary Table 12:** Gene ontology (GO) functional enrichment from large GeneMania network of ieGenes (from ieQTLs associated with infiltration phenotype at a relaxed $p < 10^{-5}$ threshold).

https://drive.google.com/drive/folders/19KvgtIgKzmTTUY5WSB3kP8ubUVtvK08T?usp=sharing
*See SupplementaryTable12.txt.*

**Supplementary Table 13:**
SNPs associated with multiple infiltration phenotypes at a relaxed $p < 10^{-5}$ threshold.

| GTEx SNP id | Tissue | Cell Type | p-value | q-value | FDR adj p-value |
|---|---|---|---|---|---|
| 7_14282722_G_A_b37 | Vagina | CD8+ T cells | $8.9 \times 10^{-6}$ | 0.17 | 0.37 |
| 7_14282722_G_A_b37 | Vagina | CD4+ T cells | $3.3 \times 10^{-6}$ | 0.23 | 0.53 |
| 18_41664299_A_G_b37 | Small_Intestine_-_Terminal_Ileum | CD8+ T cells | $9.4 \times 10^{-6}$ | 0.30 | 0.91 |
| 18_41664299_A_G_b37 | Small_Intestine_-_Terminal_Ileum | CD4+ T cells | $2.5 \times 10^{-6}$ | 0.32 | 0.54 |
| 6_35525675_A_T_b37 | Small_Intestine_-_Terminal_Ileum | CD8+ T cells | $2.3 \times 10^{-6}$ | 0.30 | 0.91 |
| 6_35525675_A_T_b37 | Small_Intestine_-_Terminal_Ileum | CD4+ T cells | $5.9 \times 10^{-8}$ | 0.06 | 0.10 |
| 4_28514830_A_C_b37 | Prostate | CD8+ T cells | $5.1 \times 10^{-6}$ | 0.06 | 0.15 |
| 4_28514830_A_C_b37 | Prostate | CD4+ T cells | $1.3 \times 10^{-6}$ | 0.42 | 0.82 |
| 12_3702789_C_A_b37 | Whole_Blood | CD8+ T cells | $8.9 \times 10^{-6}$ | 0.27 | 0.91 |
| 12_3702789_C_A_b37 | Whole_Blood | Neutrophils | $3.5 \times 10^{-7}$ | 0.23 | 0.54 |

**Supplementary Table 14:**
Target ieGenes of ieQTLs ($p < 10^{-5}$) in multiple infiltration phenotypes.

| **Tissue** | **Cell** | **Ensembl Gene ID** |
|---|---|---|
| Adipose_-_Visceral_(Omentum) | Macrophages | ENSG00000178381.7 |
| Lung | CD4+ T cells | ENSG00000178381.7 |
| Adipose_-_Visceral_(Omentum) | Macrophages | ENSG00000229043.2 |
| Lung | CD4+ T cells | ENSG00000229043.2 |
| Lung | Neutrophils | ENSG00000226686.3 |
| Esophagus_-_Mucosa | Macrophages | ENSG00000226686.3 |

**Supplementary Table 15:** List of synthetic mixes simulated for testing CIBERSORT and xCell.

https://drive.google.com/drive/folders/19KvgtIgKzmTTUY5WSB3kP8ubUVtvK08T?usp=sharing
*See SupplementaryTable15.txt.*

**Supplementary Table 16:** List of ieQTLs used to generate list of ieGenes for GeneMania.

https://drive.google.com/drive/folders/19KvgtIgKzmTTUY5WSB3kP8ubUVtvK08T?usp=sharing
*See SupplementaryTable16.txt.*

**Supplementary Table 17:** List of iQTLs used to estimate over- or under-representation of eQTLs.

https://drive.google.com/drive/folders/19KvgtIgKzmTTUY5WSB3kP8ubUVtvK08T?usp=sharing
*See SupplementaryTable17.txt.*
